# Curcumin activates distinct programming of redox metabolism towards differential regulation of gene expression mediated by Nrf1 and Nrf2

**DOI:** 10.1101/2025.07.09.663854

**Authors:** Reziyamu Wufuer, Feng Jing, Hu Shaofan, Wang Meng, Liu Keli, Zhang Yiguo

## Abstract

Curcumin (CUR), a naturally occurring phenolic small molecule, has been extensively applied in the treatment of diverse diseases for over half a century. Currently, this chemical has also been verified to exhibit a wide array of its biopharmacological activities, such as enhancing immunity, possessing antiviral, anti-cardiovascular, and anti-cancer properties. Besides, also as a nutritional supplement, CUR enables inflammatory prevention and enhance the combined efficacy of it with chemotherapy. Such characteristic of safeguarding normal organisms while treating or alleviating diseases is manifested as a common bidirectional regulation and its hormetin effect of phenolic compounds, particularly in response to stress. In this redox process, two antioxidant transcription factors Nrf1 and Nrf2 (encoded by *Nfe2l1* and *Nfe2l2*, respectively) play distinct and crucial roles. Thereby, we investigated their expression profiles of genes regulated by Nrf1 and Nrf2 in different signaling responses modulated by CUR. The resulting evidence has been provided, demonstrating that distinctive cellular metabolisms, molecular pathways, and signaling mechanisms account for Nrf1 and Nrf2, as drug targets. Both factors also play diverse roles in the anticancer effects of CUR on HepG2 cells and xenograft mice. However, the effect of CUR on xenograft tumors *in vivo* is not entirely satisfactory, although such anti-cancer effect was achieved by promoting Nrf1 expression, it appeared more reliant on Nrf2 (particularly in the absence of Nrf1). As such, we unexpectedly discovered there is not a simple regulatory relationship between CUR and Nrf2. This is supported by substantial inhibition of Nrf2 by CUR in the aberrantly proliferating *Nrf1α*^*–/–*^ cells, even albeit this chemical stimulates the expression of Nrf1 and Nrf2 in wild-type cells.

## 1. Introduction

Studies have confirmed that transcription factors can be involved in the regulation of hepatocellular carcinoma associated viral infection, tumor cell proliferation, apoptosis and metastasis, treatment and prognostic biomarkers ^[1] [2]^. Nrf1 (NF-E2-related transcription factor 1, nuclear factor 2 related transcription factor 1) and Nrf2 (NF-E2-related transcription factor 2, Both nuclear factor erythrocyte 2 associated factor 2) are functional members of the basic leucine zipper family subfamily CNC-bZIP. Several studies have shown that Nrf1 and Nrf2 play synergistic or different functions in different biological processes (proliferation, anti-inflammatory, anti-tumor, etc.) ^[3-5]^, so they are considered to be the key factors in maintaining homeostasis. With the deepening of research, Nrf1 has been found to have a stronger potential in cancer prevention and treatment, playing its unique and indispensable role. Recently, some researchers analyzed that Nrf1 is also one of the key target genes of cancer in the study of MACC1. Therefore, finding the activation of Nrf1 expression to achieve tumor defense or resistance has become a further research objective.

Natural products are the basis for the use of many drugs and are often used in the exploration of relevant active or target genes. Among them, flavonoids have a regulatory activity on numerous transcription factors not only in plants but equally in animal cells ^[6, 7]^. In related studies, it has been confirmed that low levels of CUR do not cause DNA damage, but play an antioxidant role in the carcinogenic process. High doses of CUR, on the other hand, cause oxidative stress through increased ROS production and lipid peroxidation, as well as DNA damage ^[8]^. In pancreatic cancer cells, CUR inhibits the migration and invasion of cancer cells by inhibiting ROS-mediated activation of ERK1/2 and NF-kB, resulting in decreased expression of invasion-related genes MMP2 and MMP9 ^[9]^. It is shown that CUR affects tumor development in different cancer cells by regulating oxidative stress and tumor signal transduction. Phase I and Phase II clinical studies have found that CUR has a high safety even in humans and may have anticancer therapeutic efficacy. For example, in patients with bladder cancer, cervical cancer and other tumors, increasing the CUR dose (500-8000mg /d) for 3 consecutive months can improve the precancerous lesions of patients ^[10]^. Recently, curcumin micelles have also been proposed for use as a supplement during treatment for COVID-19 patients ^[11]^. All the above results indicate that CUR has shown safe, effective and low toxicity biological characteristics in in vivo and in vitro studies. In addition, Rui Z. et al. found in CUR’s study on the cytotoxicity of inorganic arsenic that it can activate Nrf2, and the expression of Nrf1 can also be activated at a high concentration ^[12]^. However, no systematic study and explanation of Nrf1 and its mediated signaling pathway have been conducted. At present, more studies have been conducted on the relationship between the CUR analogs after structural modification and Nrf1. For example, ASC-JM17 acts on Nrf1, Nrf2 and Hsf1 to increase the expression of proteasome subunits, antioxidant enzymes and molecular companionship^[13]^. Therefore, the regulatory relationship between CUR and Nrf1 still needs to be clarified.

Recently, natural active molecules have been widely used in the study of tumor inhibition targets. Curcumin has inhibitory activity on a variety of cancer cells, and it is not only an antioxidant, but also a hormetin, which follows a bidirectional pattern in the regulation of stress response. As previous studies have shown, due to the complexity of Nrf1’s structure and function, no systematic reports on its activator (drug) have been reported. Therefore, we speculate that Nrf1 may be a key factor in the antioxidant and anti-tumor effects of curcumin. Here, Nrf1 will be used as a marker for tumor inhibition, and the effects of curcumin on Nrf1, redox and anti-tumor signal transduction will be evaluated through different genotypes of HepG2 cells and tumor xenograft models, combined with biological and omics methods, so as to clarify the targeting and anti-tumor mechanism of curcumin and Nrf1. It lays a preliminary foundation for the subsequent structural modification or the discovery of new compounds target in Nrf1.

## 2. Materials and methods

### 2.1 Chemicals and antibodies

Curcumin (Diferuloylmethane), a natural phenolic compound with formula of C_21_H_20_O_6_ and MW of 368.38 g/mol (CAS No. 458-37-7, MCE, #HY-N0005, Shanghai, China). In this study, 50.0 mg of CUR powder was completely dissolved to obtain a stock solution (DMSO, #D8371, solarbio, Beijing, China) of 200 mM CUR stored at −20°C (Follow the instructions on the purchase website: https://www.medchemexpress.cn/)” /www.medchemexpress.cn/). The antibody against Nrf1 was made in our own laboratory. These antibodies against Nrf2 (ab62352), GCLC (ab207777), GCLM (ab126704), HO-1 (ab52947), GPX1 (ab108427), XBP1 (AB109221), ATF4 (ab184909), ATF6 (ab227830), P4HB (ab137180), JUK(ab208035), p-JNK (ab124956), p-P38 (ab178876), PTEN (ab32199) and PI3K (ab302958) were obtained by Abcam (Cambridge, UK). Such antibodies against GSR (D220726), NQO1 (D26104),HK1 (D221854), ACCα (D155300), FASN (D272601), SCD1 (D162163) and CPT2 (D194980) were purchased from Sangon Biotech (Shanghai, China). Antibodies against BIP (bs-1219R), Chop (bs-20669R), p-IRE1 (bs-16698R), CDH1 (bs-10009R) and P65 (bs-0465R) were from Bioss (Beijing, China). Those antibodies against to p-eIF2α (#5199) was from CST (Boston, USA), p-PERK (sc-32577) from Santa Crus (CA, USA). The other antibodies against TXN1 (A5894), GLUT1 (A5514), GLUT4 (A11208), ATF4 (A5514), HSP60 (A5629), FGF21 (A5710), ERK (A5029), p-ERK (A5036),MEK (A5606), p-MEK (A5191) and P38 (A5017) were obtained from bimake (Texas, USA). Additional antibodies against G6PD (A1537), HK2 (A0994), LDHA (A7637), PDH (A1146) were from ABclonal (Wuhan, China). The β-actin (TA-09) was obtained from ZSGB-BIO (Beijing, China).

### 2.2 Cell line and cell culture

The human hepatocellular carcinoma HepG2 cells (i.e. Nrf1/2^+/+^ or WT) were obtained from the American Type Culture Collection (ATCC, Manassas, VA, USA). Two derived cell lines with knockout of Nrf1α^−/−^ or Nrf2^−/−^ were established in our own laboratory, and their relevant characterization had been described elsewhere by Qiu et al and Ren et al ^[14, 15]^. The fidelity of HepG2 cells had been conformed to be true by its authentication profiling and STR (short term repeat) typing map (Shanghai Biowing Applied Biotechnology Co., Ltd, Shanghai, China). All these cell lines were incubated in a 37°C with 5% CO_2_, and allowed for growth in DMEM with 25 mmol/L high glucose (Gibco, USA), 10% fetal bovine serum (Gibco, USA) and 100 units/mL penicillin-streptomycin (MACKLIN, Shanghai, China).

### 2.3 MTT assay detected cell viability

Three distinct genotypic cell lines (*WT, Nrf1α*^*–/–*^, *Nrf2*^*–/–*^) were seeded at a density of 6 × 10^3^ cells in each of 96 well plates. After the cells completely adhered, they were treated with different concentrations of CUR (0, 0.75, 1.5, 3, 6, 12, 25, 50 or 100 μM) for 24 h or 48 h. Additionally, MTT reagents (10 μL per well of 10 mg/mL stocked) was used to detect by the cell viability. Each of these experiments was repeated in five separated wells.

### 2.4 Detection of ROS and apoptosis by flow cytometry

To measure intracellular ROS levels, equal numbers (3 × 10^5^ cells in each well of 6-well plates) of experimental cells (*WT, Nrf1α*^*–/–*^, *Nrf2*^*–/–*^) were allowed for 24 h growth. After reaching 80% of their confluence, the cells were treated with 50 μM CUR for distinct time periods (i.e., 0, 12, 24 h) before being collected. The intracellular ROS were determined by flow cytometry, according to the instruction of a ROS assay kit (S0033S, Beyotime, Shanghai, China). The obtained data are presented by column charts at a ratio of Ex/Ex_*WT0*_. Next, experimental cells were treated with 50 μM CUR for 24 h and subjected to measuring cell apoptosis according to the instructions of relevant kits (C1062S, Beyotime, Shanghai, China), respectively. The resulting data are presented by column charts of a percentage or a ratio of *Ex/Ex*_*WT0*_.

### 2.5 Immunocytochemistry of γ-H2AX by confocal microscopy

Each group of cell lines (3 × 10^5^) were allowed for growth in each well of 6-well plates. After the cells were fully adhered, they were treated with 50 μM CUR for different lengths of time (0 or 24 h) and then washed with PBS, before being tested for its DNA damage. The fixed cells were blocked and then incubated with the antibody against γ-H2AX (C2035S, Beyotime, Shanghai, China), followed by DNA-staining with DAPI (4’,6-diamidino-2-phenylindole) according to the instruction of this manufacturer (I029-1-1, Nanjing Jiancheng, Nanjing, China). The immunoflorescent staining cells were subjected to confocal microscopic observation of green fluorescent γ-H2AX at λex/em= 495/519 nm and blue fluorescent DAPI at λex/em = 464/454 nm.

### 2.6 Analysis of key gene expression by real-time quantitative PCR

All experiment cells (3 × 10^5^ per well) were seeded in 6-well plates and then allowed for growth to reach 80% of their confluence, before they were treated with 50 μM CUR for 12 h or 24 h. The total RNAs were extracted and subsequently subjected to reaction with a reverse transcriptase to synthesize the first strand of cDNAs. This cDNAs were used as the template of real-time quantitative PCR (RT-qPCR) with each pair of the indicated gene primers (as listed in supplementary Table 1) to determine the mRNA expression levels of those genes in different genotypic cell lines (*WT, Nrf1α*^*–/–*^, *Nrf2*^*–/–*^). All these experiments were carried out within the Go Taq real-time PCR detection system by a CFX96 instrument. The resulting data were analyzed by the Bio-Rad CFX Manager 3.1 software and then presented graphically.

### 2.7 Analysis of key protein expression by western blotting

Experimental cell lines (*WT, Nrf1α*^*–/–*^, *Nrf2*^*–/–*^) were cultured at 6-well plates (4 × 10^5^ cells per well) and allowed for exposure into the indicated experimental settings. All those examined protein expression abundances were determined by Western blotting with their specific antibodies. Briefly, after quantitating total proteins in each of samples with the BCA protein reagent (P1513-1, ApplyGen, Beijing, China), they were separated by SDS-PAGE gels containing 8% to 12% polyacrylamide and then transferred onto the PVDF membranes. The membranes were blocked in Tris-buffered saline containing 5% non-fat dry milk at room temperature for 1 h, and incubated with each of specific primary antibodies overnight at 4°C. The antibody-blotted membranes were washed with PBST three times, before being re-incubated with the secondary antibodies at room temperatures for 2 h and then subjected to the imaging of immunoblots by Bio-Rad instrument. The intensity of immunoblotted proteins was calculated by the Quantity One software, while β-actin was used as a loading control.

### 2.8 Subcutaneous tumor xenografts in nude mice with CUR intervention

Twenty male nude mice (BALB/C^nu/nu^ aged at 6 weeks, with 16-18 g in their weights, from the HFK Bioscience, Beijing) were randomly divided into four groups: *WT_vehicle, WT_CUR, Nrf1α*^*–/–*^*_vihicle*, and *Nrf1α*^*–/–*^*_CUR*. After normal feeding for 5 d, mouse xenograft models were made by subcutaneous heterotransplantation of HepG2 (WT) and *Nrf1α*^*–/–*^ cell lines, respectively, into nude mice as described ^[16]^. Each line of experimental cells (1 × 10^7^) was allowed for its exponential growth and then suspended in 0.2 mL of serum-free DMEM, before being inoculated subcutaneously into the right upper back region of indicated mice at a single site. At the 4^nd^ day inoculated, the indicated groups of mice were subjected to intervention of CUR (35 mg/mL, by Intragastric, once a day, stopping every 5 days, for a total of 10 days). The concentration of the CUR was obtained by calculating the *IC*_*50*_ concentration in the MTT assay and based on the ratio of human to mouse surface area (a large number of studies have reported that curcumin has extremely low toxicity). Subsequently, the sizes of xenograft tumors were measured every two days and calculated by a standard formula (*i.e*., V = ab^2^/2) as shown graphically. On the 16^nd^ day after intervention by CUR, all the mice were euthanized and the tumors were removed, before being subjected to histopathological observations and other experimental analyses. Notably, all the mice were maintained under standard animal housing conditions with a 12-h dark cycle and allowed access ad libitum to sterilized water and diet. All relevant studies were accordanced with United Kingdom Animal (Scientific Procedures) Act (1986) and the guidelines of the Animal Care and Use Committees of Chongqing University (with the license SCXK-PLA-20210211), both of which were also subjected to the local ethical review in China. All relevant experimental protocols were approved by the University Laboratory Animal Welfare and Ethics Committee. Another ethical issue related to the experiment is that the tumor size of a few xenograft model mice, especially in *Nrf1α*^*–/–*^*_vehicle* group is too large to prevent certain hemorrhagic ulcers and thus shorten the feeding cycle. Overall, such related researches had indeed been conducted in accordance with approved and effective ethical regulations.

### 2.9 Histopathological examination by HE and TUNEL staining

The above-described tumor tissues of each group were fixed with 4% paraformaldehyde, and then transferred to 70% ethanol, according to the conventional protocol. Thereafter, all the individual tumor tissues were placed in the processing cassettes, dehydrated through a serial of alcohol gradient, and then embedded in paraffin wax blocks before being sectioned into a series of 5-µm-thick slides. Next, the tissue sections were dewaxed in xylene, and thereafter washed twice in anhydrous ethanol to eliminate the residual xylene, followed by rehydration in another series of gradient concentrations of ethanol with being distilled. Subsequently, they were stained with the standard HE (hematoxylin and eosin) and visualized by routine microscopy. Furthermore, these tumor tissues were prepared on a slicing machine to yield frozen sections on a glass slide, and then fixed in a constant cold box as described above. The subsequent experimental procedure was manipulated according to the instruction of an in situ Tunel (TdT-mediated dUTP Nick-End Labeling) fluorescent staining kit (G002-3-1, Nanjing Jiancheng, Nanjing, China), before being stained with DAPI. After the staining solution were removed, these tumor sections were subjected to histopathological observations under a fluorescence microscope.

### 2.10 Bioinformatic analysis of transcriptomes and metabolomes

Total RNAs were extracted from each of experimental cell lines, that had incubated with 50 μM CUR for 24 h, and then subjected to the transcriptomic sequencing by Beijing Genomics Institute (BGI, Shenzhen, China) on an Illumina HiSeq 2000 sequencing system (Illumina, San Diego, CA). All the detected mRNAs were fragmented into short fragments of approximately 200 bp. The resulting data were bioinformatically analyzed at the BGI database to determine differential expression genes (DEGs) as shown in distinct plotting forms. Thereafter, the untargeted metabolomes were employed to detect distinctions amongst three experimental cell lines (*WT, Nrf1α*^*–/–*^, *Nrf2*^*–/–*^) and between the relevant xenograft tumor tissues (*WT_vehicle, Nrf1α*^*–/–_*^*vehicle*), as conducted in three replicates of each group (Wuhan Metware Biological Co., Ltd). The obtaining data were subjected to bioinformatic analysis of differential metabolites and pathway enrichments. Subsequently, all the results from bioinformatic analyses were further validated by relevant biological experiments

### 2.11 Statistical analysis

The relevant data presented in this study are shown as fold changes (mean ± SD) relative to indicated controls, that were calculated from at least three and even five independent experiments, each of which was performed in triplicates. Statistical significances were assessed by using one-way ANOVA. Besides Student’s t-test, the Tukey’s post hoc test was also employed to determine the statistical significances for all pairwise comparisons of interest. These differences between distinct treatments were considered to be statistically significant at *p* < 0.05 (* or $) or 0.01 (** or $$).

## 3. Results

### 3.1 CUR can maintain Nrf1 protein stability through different regulation of proteasome expression

Here, we preliminatively found strong binding activity between curcumin and Nrf1 (Fig1. A) through molecular docking, the binding energy was −6.08 Kcal/mol (active-pocket 1) and −6.77 Kcal/mol (active-pocket 2). The docking sites are multiple amino acid sites in acid domain 2 (AD2), carbon terminal domain (CTD), and acid domain 1 (AD1) of Nrf1, respectively. The results of the biological experiments showed that the half-life of Nrf1 protein in HepG2 cells was significantly prolonged after CUR intervention (Fig1. B). The half-lives of Nrf1 glycoprotein (*i.e*., Nrf1-G), deglycoprotein (*i.e*., Nrf1-D), and its processed protein (*i.e*., Nrf1-P) were 0.82 h (49.2 min), 1.85 h (111 min), and 1.14 h (68.4 min), respectively (Fig1. B, b2). And the same effect, or even better, on the stability maintenance of exogenous Nrf1 proteins (Fig S1). Western-blot results showed that compared with WT cells, *Nrf2*^*–/–*^ cells were deficient in Nrf2 protein expression, while *Nrf1α*^*–/–*^ cells were deficient in Nrf1α protein expression (Fig1. C). At the same time, could induce the expression of Nrf1 protein in WT cells when CUR concentration was 6-100 μM, and significantly induced up to 9 times, and Nrf2 protein was more easily activated at 6-12 μM concentration, while other concentrations had no significant effect or inhibition on Nrf2 (Fig1. D-F). However, the induction effect of Nrf1 in *Nrf2*^*–/–*^ cells was only effective at the concentration of 6-50 μM, and showed a trend of decreasing after increasing; The expression of Nrf2 was decreased or unchanged in *Nrf1α*^*–/–*^ cells. The results showed that both Nrf1 and Nrf2 were concentration-dependent on the induction of CUR. After comprehensive comparison, the CUR of 50 μM concentration with stable Nrf1 induction effect was selected as the experimental concentration.

**Figure 1.**
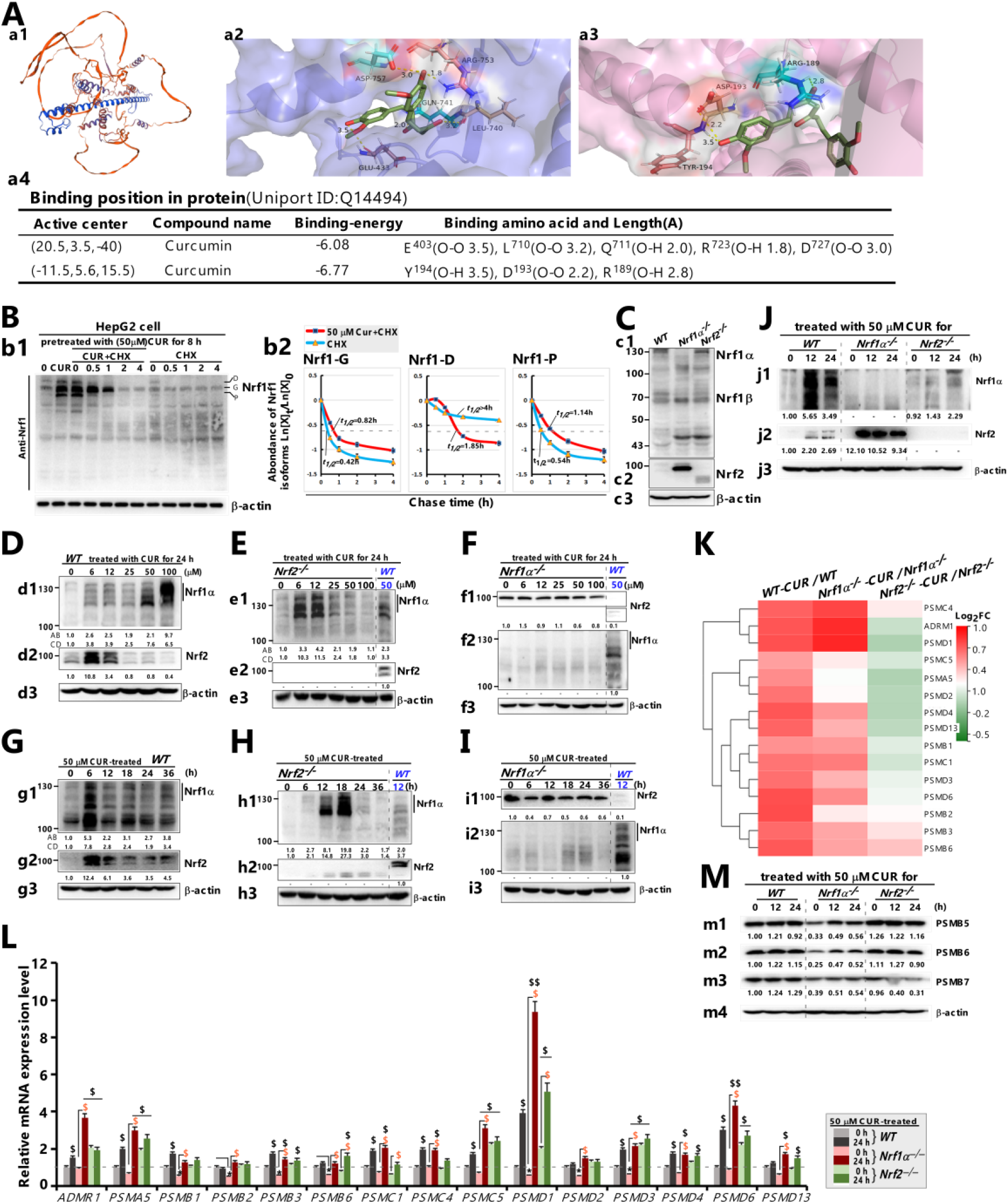
Curcumin affects the stability of Nrf1 protein through the expression level of proteasome subunits. **(A)** The 3D-Alphafold structure file ***(a1)*** of Nrf1 (long subtype TCF11: Q14494) was obtained through UNIPORT (http://www.uniprot.org/), and the active site was predicted using the DEEPSITE database (previous research has been published), and then AUTODOCK4.0 was used to connect CUR with Nrf1. The binding energies of E403, L710, Q711, R723 and D727 *aa* sites **(a2)** and Y194, D193 and R189 aa sites ***(a3)*** were −6.08 kcal/mol and −6.77 kcal/mol ***(a4)***, respectively. Note: TCF11 is a long subtype of Nrf1, containing 772 *aa*, Nrf1 is 742 *aa*, the difference between the two is that Nrf1 lacks 30 *aa* between 242 - 271 at AD1 domain in TCF11. Although the aa number is different, the docking position is the same. Similar reports have been reported in previous studies ^[17]^. **(B)** After the CUR (50 μM) interfered with the cells for 8 h, the CUR and CHX or CHX separately interfered with the cells for 0, 0.5, 1.0, 2.0, 4.0 h, respectively. Protein samples were collected and 8% SDS-PAGE adhesive was used to identify Nrf1 ***(b1)***. The results showed that the half-lives of Nrf1 glycoprotein (i.e. Nrf1-g), deglycoprotein (i.e. Nrf1-d) and its processing protein (i.e. Nrf1-p) were 0.82 h (49.2 min), 1.85 h (111 min) and 1.14 h (68.4 min), respectively ***(b2)*. (C)** Western-blot was used to detect the expression of Nrf1 and Nrf2 in different cells (*WT, Nrf1α*^*–/–*^, *Nrf2*^*–/–*^). Subsequently, **(D-F)** the expression level of Nrf1 was detected after CUR intervention at different concentrations and **(G-I)** different times. **(J)** Therefore, 50 μM CUR was used to intervene for 0, 12, and 24 h respectively to observe the expression differences among the three types of cells. **(K)** Thereafter, the total RNA was extracted from three types of cells after 50 μM CUR intervention for 0 and 24 h. After determining the concentration, transcriptome sequencing was conducted. **(M-L)** The results showed that the expression of proteasome subunit genes was differentially expressed, and the expression levels of these genes in different cells could all be affected by CUR. In addition, it is notable that the intensity of all the above-described immunoblots was also quantified by the Quantity One 4.5.2 software as shown below each of indicated protein bands, and relative their respective to β-actin after controls at WT_*t0*_. Those real-time qPCR data were shown as fold changes (mean±SD, n=3×3), which are representative of at least three independent experiments being each performed in triplicates. Significant increases ($, *p* < 0.05; $$, *p* < 0.01) and significant decreases (*\*p* < 0.05; *\*\*p* < 0.01) were statistically analyzed, when compared with their corresponding control values (measured at 0 h, i.e., *t*_*0*_), respectively. Some symbols ‘$ or *’ also indicate significant differences in the CUR intervention of *Nrf1α*^*–/–*^ or *Nrf2*^*–/–*^ cell lines, relative to their respective controls at *t*_*0*_.

After the treatment of 50 μM CUR for 0, 6, 12, 18, 24 and 36 h, Nrf1 and Nrf2 proteins were significantly expressed in WT cells (especially at 6-12 h) (Fig1. G-I). The CUR intervention for 12-18 h had the strongest effect on Nrf1 expression in *Nrf2*^*–/–*^ cells (stronger than the CUR intervention for 12 h in WT cells). Interestingly, Nrf1α was absent in *Nrf1α*^*–/–*^ cells, but the overexpressed Nrf2 decreased with the increase of CUR intervention time, but was still higher than that in WT cells after 12 h of intervention. These results indicated that the CUR could effectively induce the increase of Nrf1/2 protein level (WT and Nrf2^−/−^ cells) from 8 to 24 h, but the increase of Nrf2 in *Nrf1α*^*–/–*^ cells was significantly inhibited. In conclusion, the CUR concentration at 50 μM did significantly induce Nrf1/2 expression, but it may have a bidirectional regulatory effect on Nrf2. Therefore, (Fig1. J) we finally selected the CUR intervention concentration of 50 μM and the time of 12 and 24 h as the follow-up experimental study.

In addition, transcriptomic sequencing gene expression heat map (Fig1. K) combined with RT-qPCR results of some proteasome subunits (Fig1. L) showed that the mRNA expression of proteasome subunit genes in WT cells was significantly up-regulated after curcumin intervention, while the expression of proteasome subunit genes in *Nrf2*^*–/–*^ cell species was not significant. The protein expression results of PSMB5, PSMB6, PSMB7 and other subunits were also consistent (Fig1. M), and they were all related to the stability of Nrf1 protein.

### 3.2 CUR affects the proliferation of different genotypes HepG2 cells through differential regulation of Nrf1 and Nrf2 expression

Firstly, MTT results showed that (Fig2. A) CUR had different inhibitory effects on the growth activity of various cell lines at 0-100 μM concentration, especially on *WT* cells. The *Nrf1α*^*–/–*^ and *Nrf2*^*–/–*^ cells were only inhibited in viability when the concentration of CUR was above 20 μM. (Fig2. A, a1) Among them, the IC_50_ concentration of WT cells after 24 h of CUR intervention was 61.44 μM, that of *Nrf2*^*–/–*^ cells was 84.12 μM, while the IC_50_ value of *Nrf1α*^*–/–*^ cells was far beyond the highest concentration (100 μM) in the experiment. When the intervention time was up to 48 h, (Fig2. A, a2) the sensitivity of *Nrf1α*^*–/–*^ cells to CUR increased, and the IC_50_ concentration of WT cells was 23.59 μM, that of WT cells was 20.69 μM, and that of *Nrf2*^*–/–*^ cells was 85.04 μM. However, the viability of *Nrf1α*^*–/–*^ cells was lower than that of *Nrf2*^*–/–*^ cells regardless of the intervention time of 24 h or 48 h, while the viability of *Nrf2*^*–/–*^ cells was lower than that of *Nrf1α*^*–/–*^ cells only at higher concentrations. These results indicated that the inhibitory effect of CUR on the growth activity of HepG2 cells of the three genotypes was different, but all of them were time-dependent or concentration-dependent.

**Figure 2.**
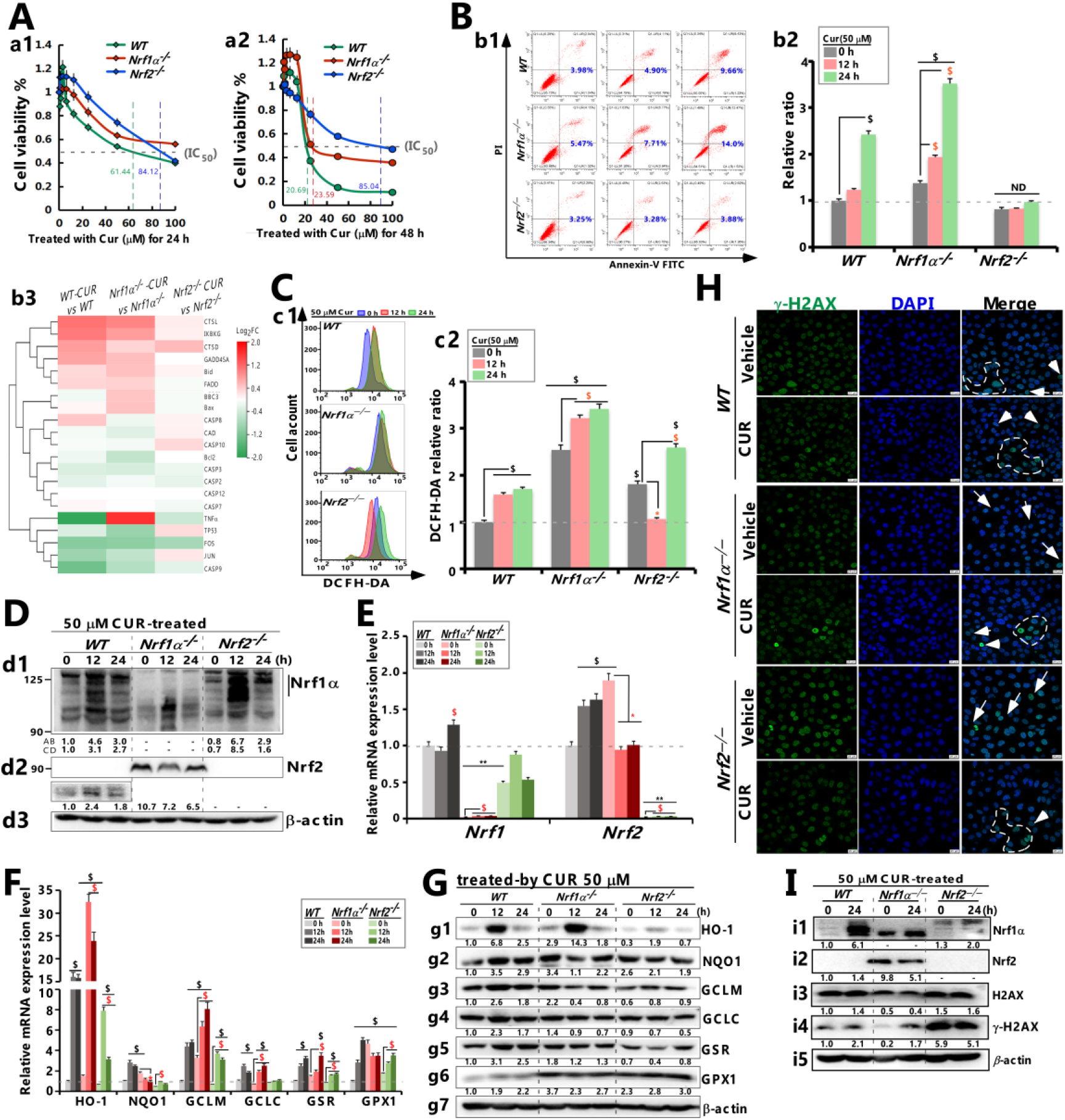
CUR differentially regulated Nrf1, Nrf2 and antioxidant gene expression, and DNA damage affected cell proliferation of different genotypes HepG2. **(A)** The cell aviability of different genotypes was measured by the enzymograph (450 nm) after CUR intervention for 24 h ***(a1)*** and 48 h ***(a2)*. (B)** Distinct apoptotic effects of CUR on different cell lines (WT, *Nrf1α*^*–/–*^ and *Nrf2*^*–/–*^) were also determined by flow cytometry, after these cell lines had been treated with 50 μM CUR for different time periods (i.e., 0, 12 and 24 h) ***(b1)***. Subsequent experimental procedure was carried out as described in the “Method and materials”. The resulting data were analyzed and also shown in the column charts ***(b2)***, which were representative of at least three independent experiments being each performed in triplicate. And differential changes of related genes in the transcriptome ***(b3)*** under the intervention of CUR. **(C)** Flow cytometry analysis of intracellular ROS levels in WT, *Nrf1α*^*–/–*^ and *Nrf2*^*–/–*^ cell lines, that had been treated with 50 μM CUR for 0, 12 or 24 h, by determining the DCFH-DA fluorescent intensity ***(c1)***. The resulting data were further analyzed by the FlowJo 7.6.1 software as shown graphical ***(c2)***, with significant increases ($$, *p* < 0.01 and $, *p* < 0.05), significant decreases (\**p* < 0.05,\*\**p* < 0.01) and no differences (ND), which were statistically determined by comparing each basal value of [*Nrf1α*^*–/–*^] and [*Nrf2*^*–/–*^] with that of WT_*t0*_. Symbols ‘$ or *’ also indicate significant differences in *Nrf1α*^*–/–*^ or *Nrf2*^*–/–*^ cells, when compared to their respective *t*_*0*_ after CUR intervention. Expression changes of Nrf1/2 **(*d1-d3*)** and **(F)** downstream antioxidant genes and proteins **(G, *g1-g7*)** after **(D-E)**CUR intervention in three genotypes of cells (WT, *Nrf1α*^*–/–*^ and *Nrf2*^*–/–*^) ; **(H)** Immunofluorescent staining of γ-H2AX that was used to detect the DNA damage of different genotypic cell lines, which was caused by CUR or vehicle (i.e., Complete medium with solvent). Subsequently, the DNA staining was conducted with DAPI. **(I)** And protein **(*i1-i5*)** levels were determined by western blotting with distinct antibodies against H2AX or γ-H2AX (i.e., phosphorylated H2AX, which serves as a typical DNA damage marker), respectively. β-actin served as a loading control. Note, the real-time qPCR data were statistically analyzed, as described in the legend for **Figure 1**.

After that, flow cytometry was used to detect apoptosis, reactive oxygen species (ROS)and DNA damage, respectively, to clarify the influence of CUR on the viability characteristics of different genotypes of HepG2 cells. The results showed (Fig2. B) that the known *Nrf1α*^*–/–*^ cell basal apoptosis level was higher than that of WT and *Nrf2*^*–/–*^ cells, while the *Nrf2*^*–/–*^ cell basal apoptosis ratio was weaker than that of WT. Compared with the basal level, the apoptosis of WT and *Nrf1α*^*–/–*^ cells after CUR intervention showed a significantly higher trend (^*$*^*p*<0.05), which was 2.5 times higher, respectively. But there was no significant difference in *Nrf2*^*–/–*^ cells compared with the basal level (*p*>0.05). Similarly, the transcriptome level of apoptosis-related genes was not significantly expressed in *Nrf2*^*–/–*^ cells (*p*>0.05), while the expression of most genes was up-regulated in *Nrf1α*^*–/–*^ cells. In addition, (Fig2. C) the level of ROS in *Nrf1α*^*–/–*^ cells was higher than that in WT (2.3 times) and *Nrf2*^*–/–*^ cells (1.8 times). After the CUR intervention, the ROS level in *Nrf2*^*–/–*^ cells decreased after 12 h of intervention (*\*p* <0.05), but the ROS level in the three types of cells was significantly increased compared with the basal expression (^*$*^*p* <0.05). At the same time, the expression levels of genes and proteins of antioxidant detoxicating enzymes down-stream of Nrf1/2 were generally highly expressed in *Nrf1α*^*–/–*^ cells, while the opposite was true in *Nrf2*^*–/–*^ cells. The CUR intervention increased the expression of these genes to a certain extent (Fig2. D-G). Although CUR has been reported to have antioxidant capacity, it still does not reduce ROS accumulation in different genotypes, and apoptosis is also associated with it, suggesting that CUR may also trigger other cell death mechanisms through Nrf1/2 in addition to the regulation of different free radicals in ROS production and elimination (antioxidant detoxification enzyme).

Related studies have reported that highly activated ROS can cause mitochondrial and nuclear DNA damage, so the effect of CUR on three types of intracellular DNA damage was further examined. The immunofluorescence observation (Fig2. H) showed that the CUR had inconsistent DNA damage induction effect on the three types of cells. Although WT cells showed no significant DNA damage before and after CUR intervention, their volume was generally reduced; *Nrf1α*^*–/–*^ cells showed DNA damage after the addition of CUR, while *Nrf2*^*–/–*^ cells showed no significant change. In addition, Western-blot results (Fig2. I) showed that although the basal expression of γ-H2AX was higher in *Nrf2*^*–/–*^ cells, it was significantly up-regulated in WT and *Nrf1α*^*–/–*^ cells after CUR intervention (especially in *Nrf1α*^*–/–*^ cells). The above results indicated that, although Nrf1 was weakly expressed in *Nrf2*^*–/–*^ cells, it still played a role in cell protection (it is worth mentioning that *Nrf2*^*–/–*^ cells lost the tumorigenicity), while CUR further promoted the expression of γ-H2AX in *Nrf1α*^*–/–*^ cells, weakened Nrf2, and induced DNA damage.

### 3.3 CUR affects oxidative stress progression in HepG2 cells through Nrf1/2 differential regulation of antioxidant-related gene expression

The regulation of ROS in cells is mainly carried out through the generation and elimination of ROS. Previous studies have described that Nrf1 and Nrf2 can affect the intracellular antioxidant process by binding to the *ARE* sites of redox related genes ^[4]^. The ROS generation/elimination system, the GSH synthesis system (the conversion process between GSH and GSSG), the NADPH synthesis system (related to reprogramming of redox metabolism), the TXN system (oxidized and reduced transformation), and the metal (iron) homeostasis system (mainly the transformation between Fe^+3^ and Fe^+2^) can be regarded as the key to the cellular antioxidant system Influencing factors.

Firstly, transcriptomic results showed (Fig3. A), the expression of *FTL* and *GPX2* in *Nrf1α*^*–/–*^ cells and *SESN3* and *SLC7A11* in *Nrf2*^*–/–*^ cells were significantly increased compared with the basal level except for some genes related to iron homeostasia, GSH system and NADPH synthesis in WT cells (^*$*^*P*<0.05; ^*$$*^*P*<0.01), the differential expression of other genes decreased or had no significant difference. After, combined with the results of RT-qPCR (Fig3. B-F), it was found that in WT cells, GSH antioxidant system genes (*GCLC, GCLM, GSR, GLS, SLC7A11*), iron homeostatin genes (*HO-1, SESN1, SESN2, SESN3*), TXN antioxidant genes (*TXN1, TXN2, SRXN1*), NADPH synthesis-related genes (*IDH1, IDH2, ME1, TIGAR*) The mRNA levels of ROS generation/eliminating genes (*CAT, PRDX1, SOD1, SOD2*) were significantly increased after CUR intervention (^*$*^*p*<0.05). When CUR intervened, GSH antioxidant system (e.g., *GCLC, GCLM, GSR*), iron homeostasis genes (e.g., *HO-1, SESN1*), TXN system genes (e.g., *TXN1, SREN1*), NADPH synthesis genes (*IDH1, IDH2, TIGAR*) and ROS producing genes (e.g., *CAT, PRDX1, PRDX3, GPX2* and *SOD1*) in *Nrf1α*^*–/–*^ cells, the mRNA expression levels of were significantly increased (^*$*^*p* <0.05; ^*$$*^*p* <0.01). And the mRNA levels of other genes such as *SOD2, G6PD, SLC7A11* and *ME1* were significantly decreased (*\*p*<0.05). However, the mRNA expressions of *SOD1, IDH1, IDH2, GCLM, SLC7A11, SESN3* and other genes were only up-regulated in *Nrf2*^*–/–*^ cells after CUR intervention (^*$*^*p*<0.05). Although CUR induced genes promoting the generation of ROS (such as ·O^-2^ and H_2_O_2_) genes SOD1/2 and IDH1/2, it also induced the expression of antioxidant genes with stronger reducing ability such as *CAT* and *GPX2*. At the same time, the mRNA levels of *GCLC, GCLM, GSL, PRDX, SESN, G6PD* and *TIGAR* were significantly promoted (^*$*^*p*<0.05), so as to weaken the production of ROS (H_2_O_2_, ·OH, ·O^-2^, etc). In addition, according to the protein expression of some genes (Fig3. G, g1-g8), it was found that CUR had the same inducing effect on the G6PD, HO-1, GCLC, TXN1 and SOD2 in WT cells. But, in *Nrf1α*^*–/–*^ and *Nrf2*^*–/–*^ cells, the protein levels of other genes were not significantly affected except for the decreased expression of HO-1 (in *Nrf1α*^*–/–*^) and TXN1 (in *Nrf2*^*–/–*^). The results showed that CUR significantly regulated the mRNA levels of antioxidant system genes in different genotypes. In general, Nrf1 and Nrf2 made different contributions in the process of exerting redox-regulatory functions on CUR, which may be the main reason why ROS level in *WT* and *Nrf1α*^*–/–*^ cells increased instead of decreasing after CUR intervention.

**Figure 3.**
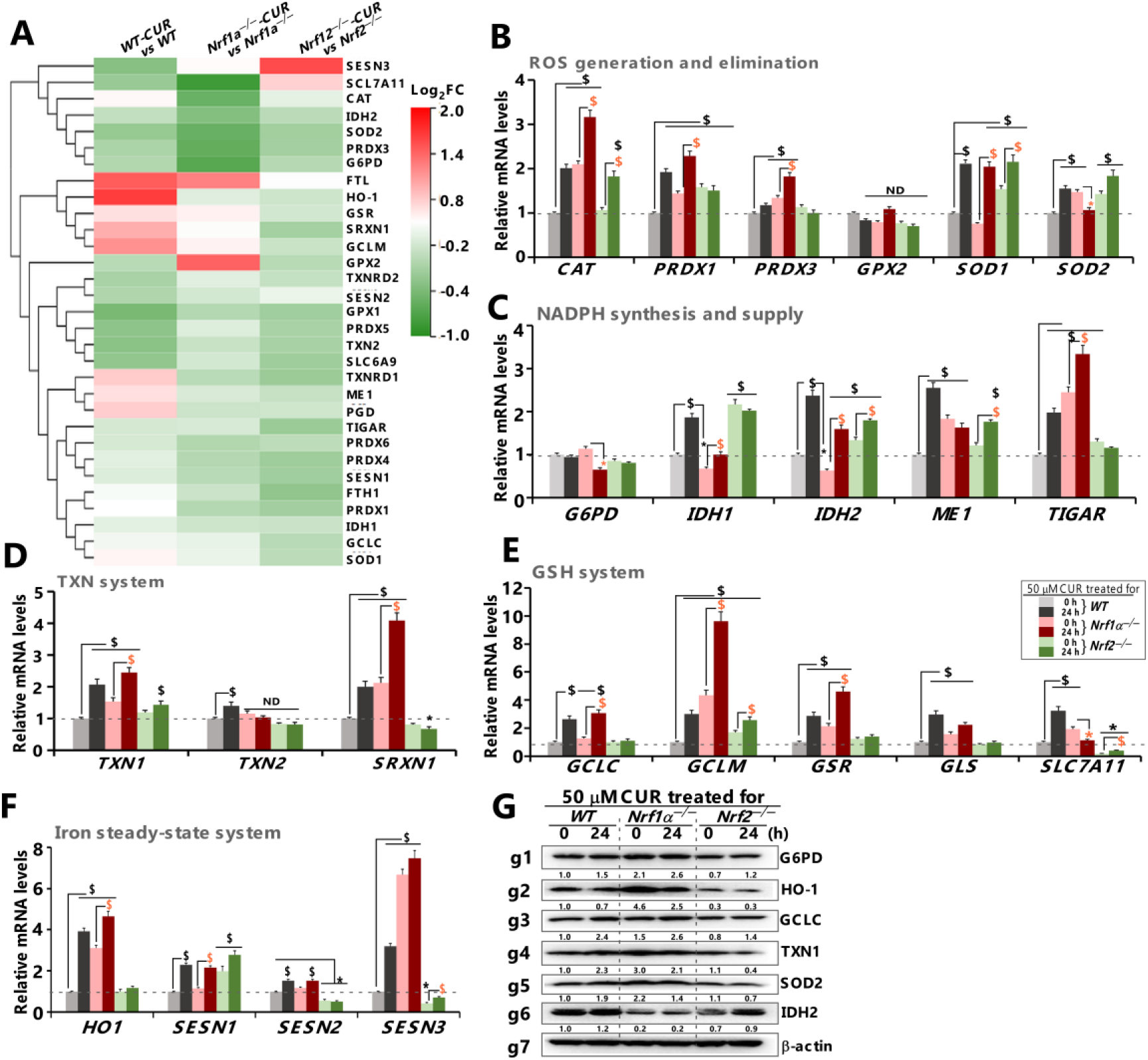
Discrete induction of Nrf1/2-mediated redox-related gene expression in different cellular responses to CUR. **(A)** The heat-map shows differential expression levels of those antioxidant and detoxification genes determined by transcriptome sequencing of WT, *Nrf1α*^*–/–*^ and *Nrf2*^*–/–*^ cell lines that had been intervened with 50 μM CUR for 0 or 24 h. **(B-F)** Real-time qPCR analysis of basal and CUR-stimulated expression levels of those genes responsible for redox homeostasis, e.g., **(B)** balancing the ROS production and elimination (*CAT, PRDX1, PRDX3, GPX2, SOD1, SOD2*), **(C)** NADPH synthesis genes (*G6PD, IDH1, IDH2, ME1, TIGAR*), **(D)** GSH biosynthesis and transport (*GCLC, GCLM, GSR, GLS, SLC7A11*), **(E)** thioredoxin (TXN) antioxidant system (*TXN1, TXN2, SRXN1*) and **(F)** Iron homeostasis (*HO-1, SESN1, SESN2, SESN3*) in WT, *Nrf1α*^*–/–*^ and *Nrf2*^*–/–*^ cell lines that hat had been intervened with 50 μM CUR for 24 h or not. The resulting data were shown as fold changes (mean±SD, n=3×3), which are representative of at least three independent experiments being each performed in triplicates, with significant increases ($$, *p* < 0.01 and $, *p* < 0.05), significant decreases (*\*p*<0.05, *\*\*p* < 0.01) and no significant differences (ND), as compared each basal value of [*Nrf1α*^*–/–*^] and [*Nrf2*^*–/–*^] with that of WT_*t0*_. Symbols ‘$ or *’ also indicate significant differences in *Nrf1α*^*–/–*^ or *Nrf2*^*–/–*^ cells, when compared to their respective *t*_*0*_ after CUR intervention. **(G)** Western blotting analysis of the indicated proteins, including G6PD, HO-1, GCLC, TXN1, SOD2, IDH2 and β-actin **(*g1-g7*)** in WT, *Nrf1α*^*–/–*^ and *Nrf2*^*–/–*^ cell lines that had been intervened with 50 μM CUR for 24 h or not. The intensity of these immunoblots was also quantified by the Quantity One 4.5.2 software and showed under those indicated protein bands. β-actin served as a loading control, and the experimental results were presented by calculating [(Ex/β-actin)/WT_*t0*_].

### 3.4 Distinctive requirements of Nrf1 and Nrf2 for different genotype cellular signaling responses to CUR

Previous studies have confirmed that Nrf1 or Nrf2 deficiency can induce stress of varied degrees in ER and mitochondria in cells ^[18]^. Therefore, we further examined relevant indicators to clarify the difference in the influence of CUR on endoplasmic reticulum (ER), mitochondrial stress and UPR phenomena through the mediation of Nrf1 or Nrf2. Transcriptomics showed (Fig4. A) that after CUR intervention, the differential expression of ER and Mito stress-related genes in the three genotypes cells were generally down-regulated or had no difference. Through the detection of the mRNA levels of related genes (Fig4. B), it was found that in WT cells, the mRNA expressions of genes such as *PDI, PERK, eIF2α, ATF4, IRE1, ATF6, CHOP*, and *FGF21* were significantly increased (^*$*^*P* < 0.05; ^*$$*^*P* < 0.01), while that of *BIP* was significantly decreased (*\*P* < 0.05), and there were no significant differences in others. In *Nrf1α*^*–/–*^ cells, the basal level of genes such as *PDI, BIP, PERK, eIF2α, ATF4, IRE1, XBP1, ATF6, CHOP, FGF21* and *HSP60* were all lower than those in WT cells. However, after CUR intervention, the mRNA levels of these genes were promoted. At the same time, the above genes were also activated in *Nrf2*^*–/–*^ cells with significant differences (^*$*^*P*< 0.05; ^*$$*^*P*< 0.01). Interestingly, after 24 h of CUR intervention, some genes (e.g., *PDI, BIP, PERK, eIF2α, XBP1*) returned to normal levels. In addition, in the results of protein level detection (Fig4. C), the protein expression trend of related genes in WT cells was consistent with the mRNA results. such as PDI, BIP, (p-) PERK, P-EIF2α, IRE1, ATF4, XBP1, ATF6, FGF21 and other proteins were significantly up-regulated after CUR intervention. After CUR intervention in *Nrf1α*^*–/–*^ cells, only the expressions of BIP, p-eIF2α, ATF4 and CHOP proteins were up-regulated, and the phosphorylation level of eIF2α and the expressions of PDI and FGF21 proteins were down-regulated. In *Nrf2*^*–/–*^ cells, CUR promoted the levels of PDI, (P-)PERK, ATF4, P-IRE1, XBP1, FGF21 and other proteins, but the phosphorylation level of eIF2α was decreased. These results suggest that CUR is more likely to participate in the ER stress signaling mediated by PDI-BIP-IRE1/PERK/ATF6 in WT and *Nrf2*^*–/–*^ cells, and affect the intracellular UPR process. It is mentioning that when Nrf1 is absent, CUR may be more likely to participate in oxidative stress and UPR processes by regulating mRNA levels of related genes (compared with protein level regulation).

**Figure 4.**
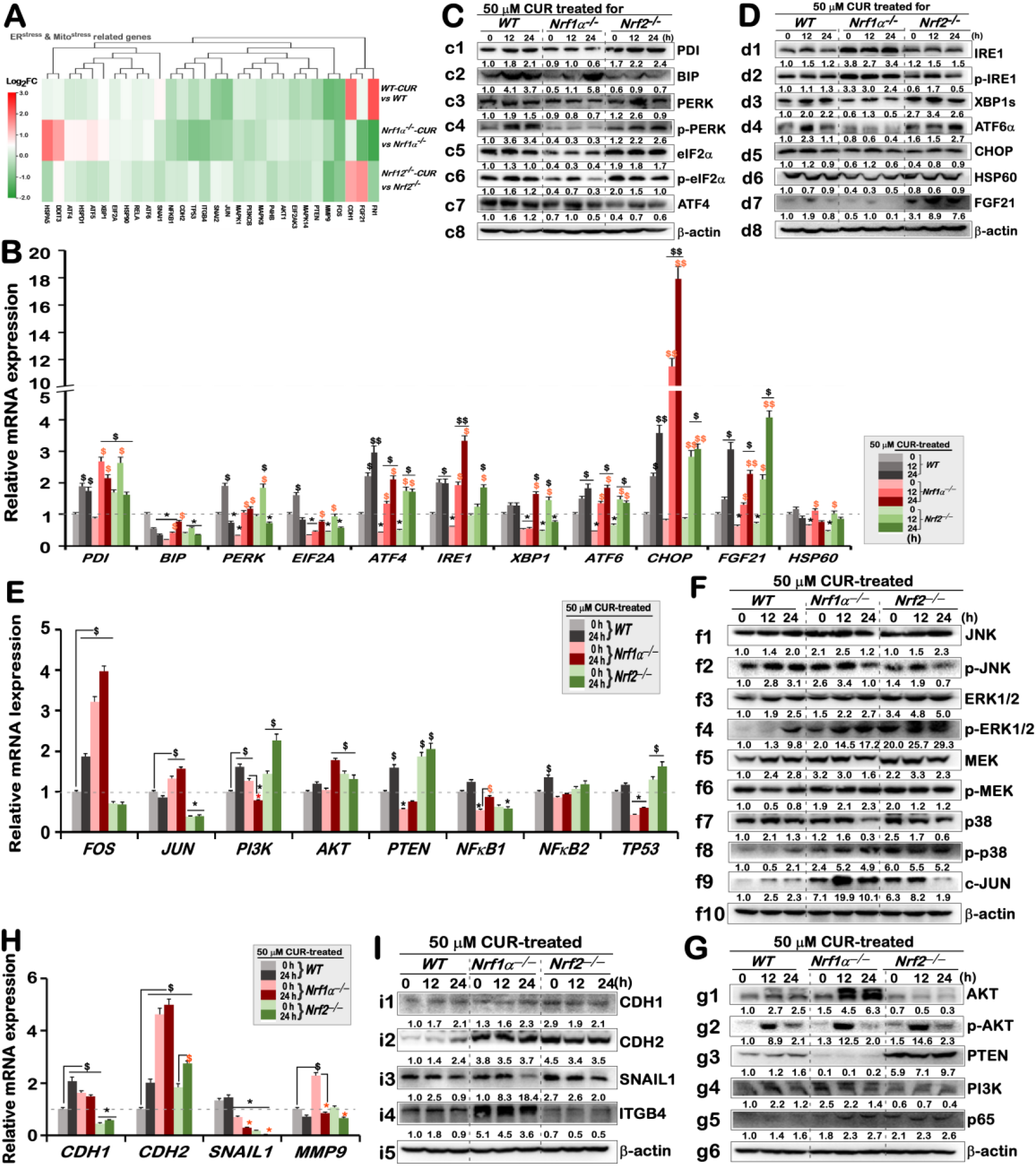
CUR regulates oxidative stress and tumor signaling pathway genes through Nrf1/2 differences to affect tumor cell fate. **(A)** The heat-map shows changes of those differential expression genes required for multi-hierarchical signaling networks (e.g., UPR, MAPK, PI3K/AKT-PTEN and EMT pathways), which were determined by transcriptome sequencing of three genotypes cell (i.e., WT, *Nrf1α*^*–/–*^ and *Nrf2*^*–/–*^ ) that had been intervened with 50 μM CUR. **(B)** Real-time qPCR analysis of the mRNA expression levels of those genes responsible for the ER and/or mitochondrial stresses (*PDI, BIP, PERK, eIF2α, ATF4, IRE1α, XBP1, ATF6, CHOP, HSP60* and *FGF21*) in three genotypes cells that had been treated with 50 μM CUR for 0, 12 or 24 h. **(C,D)** Western blotting analysis of those expression abundances of indicated proteins involved in the ER and mitochondrial stress responses after CUR stimulation of different genotypes cell lines for 0, 12 and 24 h. β-actin served as a loading control. **(E)** Real-time qPCR analysis of basal and CUR-stimulated expression levels of those genes responsible for MAPK, PTEN-PI3K-AKT pathways in three genotypes cell lines that had been intervened by CUR for 0 or 24 h. **(F, G)** Abundances of those indicated proteins involved in the MAPK, PTEN-PI3K-AKT signaling pathways were determined by Western blotting of three genotypes cell lines that had been intervened by CUR for 0, 12 or 24 h. **(H)** Real-time qPCR analysis of basal and CUR-stimulated expression levels of those genes responsible for EMT pathways in three genotypes cell lines that had been intervened by CUR for 0 or 24 h. **(I)** Abundances of those indicated proteins involved in the EMT signaling pathways were determined by Western blotting of three genotypes cell lines that had been intervened by CUR for 0, 12 or 24 h. Note: The intensity of those immunoblots was quantified by the Quantity One 4.5.2 software and also showed below the indicated protein bands (calculated by [(*Ex/β-actin*)/WT_*t0*_]). The RT-qPCR resulting data were shown as fold changes (mean±SD, n=3×3), which are representative of at least three independent experiments being each performed in triplicates. Significant increases ($, *p*<0.05; $$, *p*<0.01) and significant decreases (*\*p* < 0.05; *\*\*p* < 0.01) were statistically determined, when compared with WT controls (measured at 0 h), respectively. Additional “$ or *” symbols indicate significant differences in *Nrf1α*^*–/–*^ or *Nrf2*^*–/–*^ cell lines that had been intervened by CUR as compared with their respective *t*_*0*_ values after intervention.

Our previous studies revealed that Nrf1 deletion generates abnormal ROS aggregation, activates MAPK and PI3K/AKT signaling pathways, and thus triggers the malignant proliferation of *Nrf1α*^*–/–*^ cells ^[18, 19]^. Here, (Fig4. E-G) we found that after CUR intervention, the protein expressions of JNK, ERK1/2, MEK, P38 and other genes were up-regulated in WT cells, and the protein phosphorylation levels of JNK, ERK1/2, and P38 were enhanced. After CUR intervention, the expression of (p-) ERK1/2, P-MEK and P-P38 proteins in *Nrf1α*^*–/–*^ cells was increased, but the expression of (p-) JNK, MEK and p38 proteins was decreased. In *Nrf2*^*–/–*^ cells, CUR also promoted the expression of JNK and p-ERK1/2, and decreased the expression of p-MEK and p38. However, the expression of the transcription factor c-JUN downstream of Nrf1/2 was significantly up-regulated in WT and *Nrf1α*^*–/–*^ cells, but decreased in *Nrf2*^*–/–*^ cells. Meanwhile, CUR promoted the protein expression of (p-) AKT, PI3K and PTEN in WT cells. In *Nrf2*^*–/–*^ cells, AKT was decreased but its phosphorylation level was increased, and the expression of PTEN protein was also enhanced, while the expression of PI3K and p65 was not difference. CUR inhibited the expression of PI3K in *Nrf1α*^*–/–*^ cells, while promoting the expressions of (p-)AKT and p65, with no change in the expression of PTEN. The results showed that CUR could induce the expression and phosphorylation of most MAPK and PI3K/AKT signaling pathway genes in WT cells, while it was more likely to promote the phosphorylation of a few proteins in Nrf1 or Nrf2 knockout cells.

In tumors diseases, the epithelial-mesenchymal transition (EMT) is usually closely associated with tumorigenesis, invasion, metastasis, and therapeutic resistance. As can be seen from the mRNA results (Fig4. H), the basal expressions of *CDH1, CDH2*, and *MMP9* genes are relatively high in *Nrf1α*^*–/–*^ cells, indicating that after the knockout of Nrf1, it is more likely to activate the mRNA expression of genes related to migration. After the intervention of CUR, the expressions of *MMP9* and *SNAI1* were downregulated in all three types of cells. The results of Western-blot also showed (Fig4. I) that the expressions of the EMT marker protein SNAIL1 and the epithelial-mesenchymal protein ITGB4 were downregulated in the three types of cells (especially in *Nrf1α*^*–/–*^ cells). The results indicated that CUR could affect the migration process of tumor cells by inhibiting the mRNA levels of SNAIL1 and MMP9 genes and the protein expressions of SNAIL1 and ITGB4.

### 3.5 Reprogramming of the cellular metabolome by Nrf1α^−/−^ or Nrf2^−/−^ and influence on the metabolic response to CUR

Previous studies by our group found that the impairment of mitochondrial function in Nrf1 knockout cells led to the increase of ROS, the weakening of protein stability, and the enhancement of lipid metabolism, glucose metabolism, and amino acid metabolism, indicating that Nrf1 is a key gene in tumor metabolic reprogramming ^[18, 20]^. Here, we found that CUR is involved in Nrf1/2-mediated regulation of gene expression related to tumorigenosis and development, and these signal-transduction is also closely related to metabolic activities, so we conducted follow-up tests on metabolism-related indicators. According to the results (Fig5. A-C), the basal value of glucose, ATP and NADP^+^/NADPH in *Nrf1α*^*–/–*^ cells were much higher than those in WT and *Nrf2*^*–/–*^ cells, indicating that the depletion of Nrf1 caused A high demand for different nutrients in cells. After CUR intervention, glucose production in *Nrf1α*^*–/–*^ cells was significantly inhibited (*\*p*<0.05), but the inhibitory effects on ATP and NADP^+^/NADPH were not significant. Similarly, CUR just significantly down-regulated ATP production in WT cells (*\*p* <0.05), although it did not significantly decreased glucose, NADP^+^/NADPH production both in WT and *Nrf2*^*–/–*^ cells.

**Figure 5.**
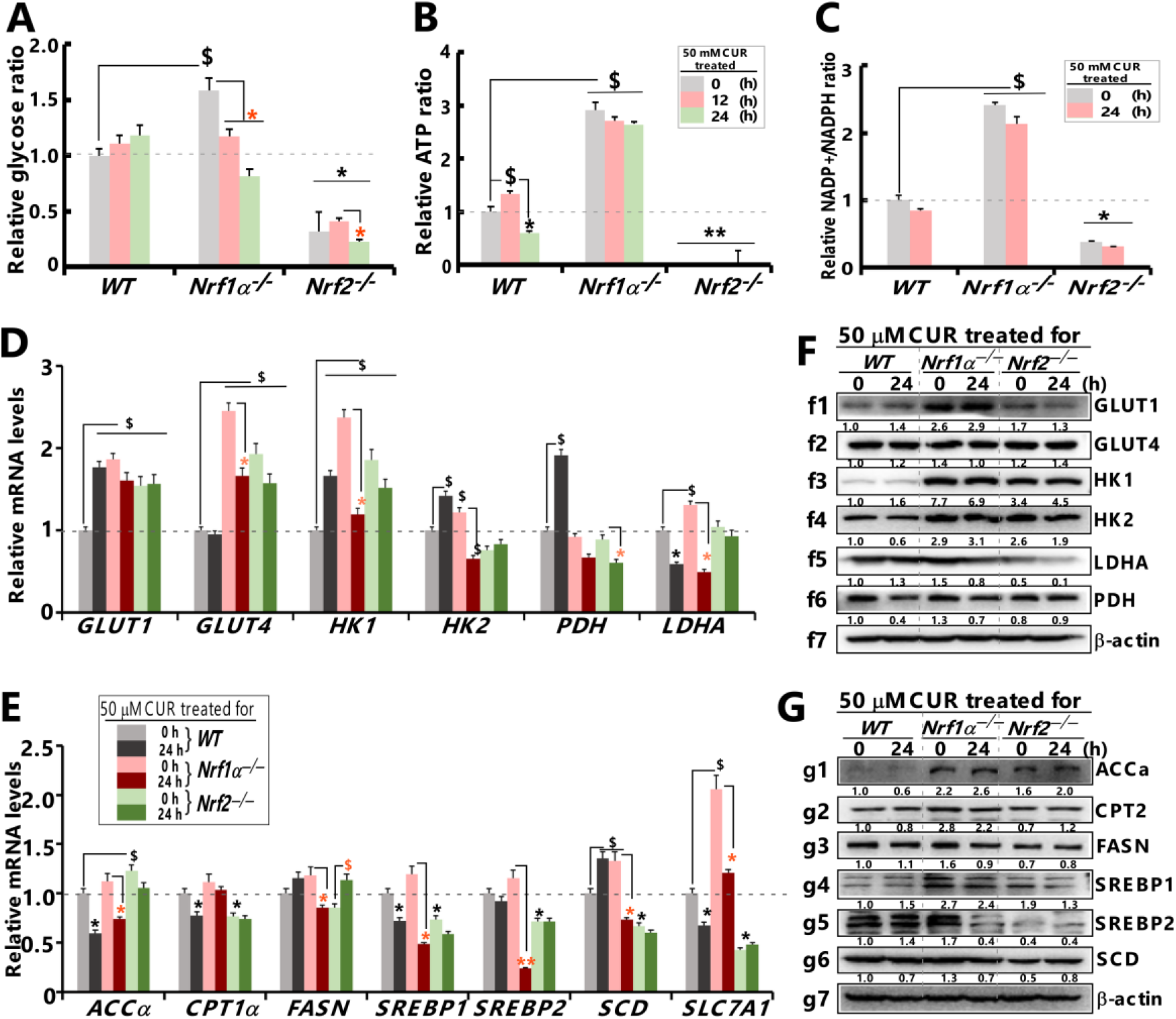
Reprogramming of the cellular metabolome by Nrf1α^−/−^ or Nrf2^−/−^ and influence on the metabolic response to CUR. **(A-C)** The content of intracellular glucose **(A)**, the ratio of ATP **(B)** and NADP+/NADPH **(C)** were determined in three genotypes cells (i.e., WT, *Nrf1α*^*–/–*^ and *Nrf2*^*–/–*^), that had been intervened with 50 μM CUR for 0, 12 or 24 h. Each group of samples was repeated in at least three composite wells. The measured absorbance (OD) values were then statistically analyzed and graphically shown as mean±SD (n=3×3), along with significant increases ($, *p*< 0.05) and significant decreases (*\*p*< 0.05; *\*\*p*< 0.01), were statistically analyzed when compared with WTt0, respectively. Additional symbols “$ or *” indicate significant differences in three genotypes cells compared to their respective *t*_*0*_ value measured after CUR intervention. **(D, E)** Real-time qPCR analysis of basal and CUR-stimulated expression levels of those genes responsible for glucose **(D)** and lipid **(E)** metabolism in three genotypes cell lines that had been intervened. **(F,G)** Abundances of those indicated proteins involved in the glucose **(F)** and lipid **(G)** metabolism were determined by Western blotting of three genotypes cell lines that had been intervened by CUR for 0, 12 or 24 h. Note: The intensity of those immunoblots was quantified by the Quantity One 4.5.2 software and also showed below the indicated protein bands (calculated by [(*Ex/β-actin*)/WT_*t0*_]). The RT-qPCR resulting data were shown as fold changes (mean±SD, n=3×3), which are representative of at least three independent experiments being each performed in triplicates. Significant increases ($, *p*<0.05; $$, *p*<0.01) and significant decreases (*\*p* < 0.05; *\*\*p* < 0.01) were statistically determined, when compared with WT controls (measured at 0 h), respectively.

RT-qPCR results showed (Fig5. D) that, the glycolysis and lipid synthesis-related genes *LDHA, ACCα, CPT1* and *SREBP1* were down-regulated, while *GLUT1, HK1, HK2* and *PDH* levels were significantly up-regulated (^$^*p*<0.05, ^$$^*p*<0.01) after CUR intervention. (Fig5. E) But there were no significant changes in *GLUT1* and *CPT1* in *Nrf1α*^*–/–*^ cells, but the expressions of other genes were significantly down-regulated (\**p*<0.05, \*\**p*<0.01). However, there was no significant difference in the above genes in *Nrf2*^*–/–*^ cells. In addition, (Fig5. F) CUR could inhibit the protein expression of HK2, PDH, CPT, SCD and others in WT cells, and promote the expression of HK1 and LDHA proteins, while the other protein levels had no difference. GLUT4, HK2, LDHA, SREBP1 and other proteins were decreased in *Nrf2*^*–/–*^ cells, but there were no changes in other proteins. (Fig5. G) CUR inhibited the expression levels of LDHA, PDH, CPT2, FASN, SREBP1, SREBP2 and SCD in *Nrf1α*^*–/–*^ cells. According to the results, CUR showed more significant regulation of mRNA and protein related to glycolysis and fatty acid synthesis in *Nrf1α*^*–/–*^ cells (which was consistent with the results of lipid drop-formation experiments (Fig S3. A)). These results indicate that CUR regulates genes related to glucose and lipid metabolism through differential expression of Nrf1/2, thereby alleviating metabolic abnormalities in tumor cells.

### 3.6 CUR weakly inhibited xenograft tumor but still affected metabolic process

Herein, we continued to examine the regulatory role of CUR on Nrf1/2 and its effect on tumors *in vivo*. It was confirmed that *Nrf1α*^*–/–*^ _*vehicle* group were rapidly grown, with approximately 4.2 times larger in size than that of *WT_vehicle* tumor. CUR intervention was administrted on the 4^th^ day of subcutaneous tumor formation and then its inhibitory effect of this intervention on xenograft tumors was seemingly observed to be roughly affected by the absence of Nrf1α (in *Nrf1α*^*–/–*^ _*CUR* group), but no change in *WT_CUR* group. (Fig6. A) (FigS4. A). Further histopathological and immunohistochemical examination (HE and TNUEL staining) revealed (Fig6. B-C) that the increased number of positive cells in tumor tissue of *Nrf1α*^*–/–*^_*vehicle*, and the degree of malignity was still higher than that of *WT_vehicle*, which was consistent with the cell experiment. (Fig6. D-F) In addition, the production of glucose, ATP and NADP^+^/NADPH in *Nrf1α*^*–/–*^_*vehicle* tumor tissue was also much higher than that of *WT_vehicle*, indicating that the malignant growth of *Nrf1α*^*–/–*^_*vehicle* was closely related to substance metabolism. After CUR treatment, both glucose and ATP production decreased in *Nrf1α*^*–/–*^_*CUR*, but completely reversed in *Nrf1α*^*–/–*^_*CUR* tissue. This might be caused by the difference between the diverse in vivo systems and the single cellular growth environment. Alternatively, it could be that CUR has triggered changes in other signaling molecules, which have accelerated the uptake of nutrients and energy. Therefore, in the following experiment, we carried out untargeted-metabolomics detection on the tumor tissues before and after the CUR intervention.

**Figure 6.**
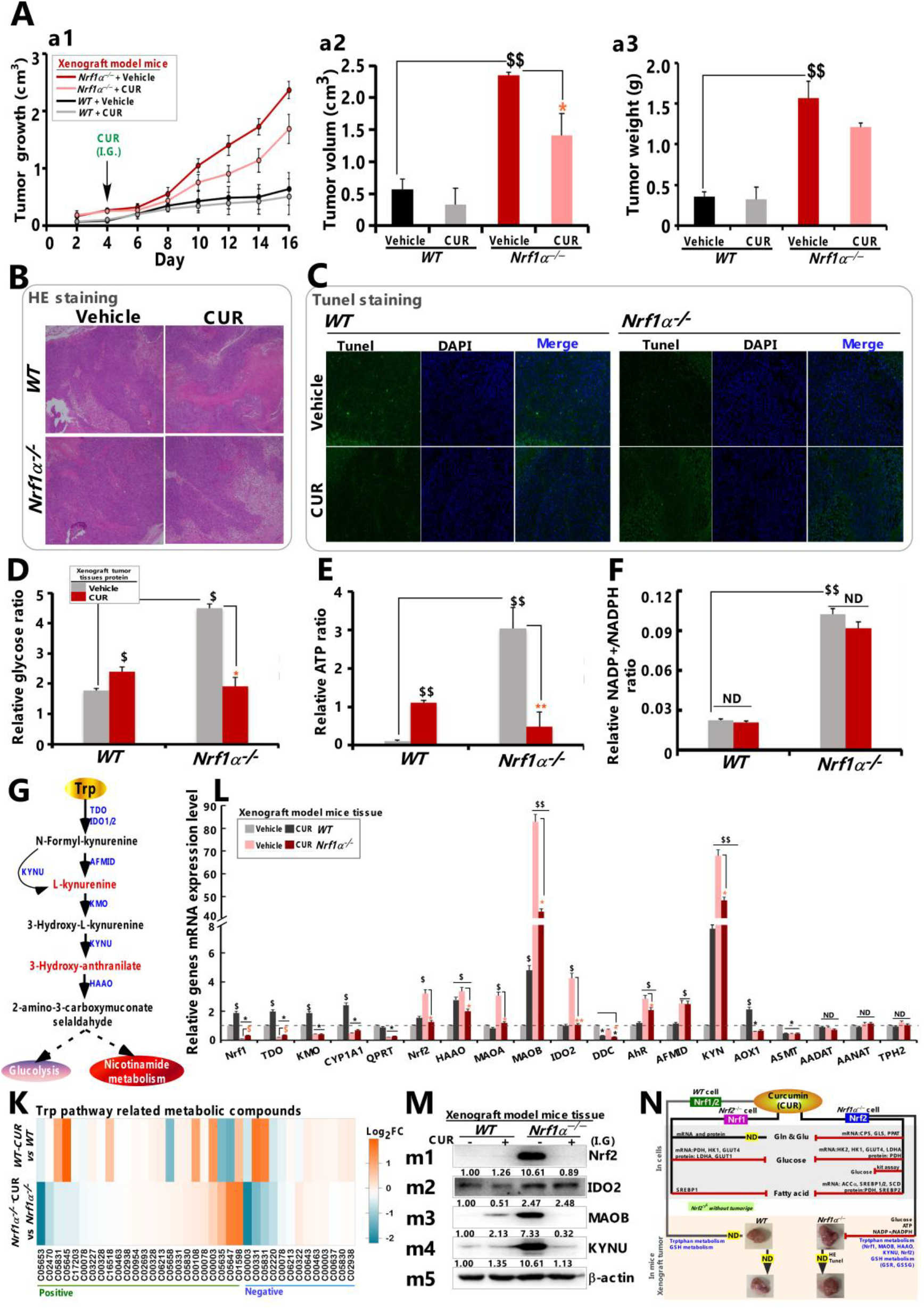
CUR weakly inhibited xenograft tumor but still affected metabolic process. **(A)** The in *vivo* malgrowth curve ***(a1)*** of xenograft tumors in distinct groups of nude mice, that were subcutaneously injected with human WT or *Nrf1α*^*–/–*^ hepatoma cell lines, followed by intervention with *CUR* (I.G., 35 mg/mL) or *vehicle* (I.G., 0.9% physiological saline). And the volumes ***(a2)*** and weight ***(a3)*** of the above xenograft tumors were determined as shown graphically. **(B)** The model tumor tissues were subjected to the histopathological examination by routine HE staining, and then relevant images were obtained from microscopy. **(C)** The staining of TUNEL was subjected to evaluating cell apoptosis in xenograft tumor tissues. **(D-F)** The content of glycose **(D)**, the ratio of ATP **(E)** and NADP^+^/NADPH **(F)** within tumor tissues were detected, which were performed at least in triplicates. The obtained absorbance (OD) values are statistically determined and shown graphically, with significant increases ($, *p*<0.05) and significant decreases (*\*p* < 0.05), relative to the control value of each group as indicated. Additional symbols “$ or *” indicate significant differences in tumor tissues compared to their respective t0 value measured after CUR intervention. **(G)** Schematic diagram shows tryptophan metabolic pathway (adapted from the online KEGG pathway database). The “**black**” means metabolic compounds, “**red**” represents key metabolites, and “**blue**” represents different metabolic enzymes. **(H)** CUR differential expression heat map of related metabolites in tryptophan metabolism in different tumor tissues (i.e., l Log_2_FC l≥1). **(L)** Real-time qPCR analysis was subjected to detecting the effects of CUR treatment on transcriptional expression of those key enzymes (e.g., *Nrf1, TDO, KMO, CYP1A1, QPRT, Nrf2, HAAO, DDC, AhR, AFMID, KYN, AOX1, ASMT, AADAT, AANAT, TPH2*) in tryptophan metabolism pathway. The resulting data were shown as fold changes (mean±SD, n =3×3), which are representative of at least three independent experiments being each performed in triplicates. Significant increases ($, *p* < 0.05; $$, *p* < 0.01) and significant decreases (*\*p* < 0.05), were statistically analyzed when compared to *WT_vehicle*, respectively. Additional symbols “$ or *” indicate significant differences in *Nrf1α*^*–/–*^ tumors as compared to their vehicle controls. “ND” means no significant different. **(M)** Western blotting analysis was subjected to detecting the protein expression of Nrf2, IDO2, MAOB and KYNU in different tumor tissues. The intensity of those immunoblots was also quantified by the Quantity One 4.5.2 software, and showed below the indicated protein bands. **(K)** Overview of the effects of CUR on different mechanisms through Nrf1/2 differential expression at the cellular and animal levels.

In our previous study, the differential pathway between *Nrf1α*^*–/–*^_*vehicle* and *WT_vehicle* tumor tissues was preliminarily identified as tryptophan metabolism ^[17]^ (without treated by CUR) (Fig6. G). Therefore, we first observed whether CUR affects the expression of related metabolic enzymes through the TRP metabolic pathway. The result shows that (Fig6. K, L), after treatment with CUR, there were generally no differences in the metabolites in this pathway, but there was differential regulation of the 3-HA level and the mRNA expression of some metabolic enzymes (i.e., *IDO2, MAOA, HAAO, DDC*). According to the report, the TRP metabolites L-Kynurenine (Kyn), 5-HA, and 3-HA are considered to provide an escape guarantee for ferroptosis in tumor cells ^[21]^. The expressions of the lipid metabolism-related enzymes KYNU and HAAO were indeed highly expressed in the *Nrf1α*^*–/–*^_*vehicle* tumor tissues, and the intervention of CUR could significantly inhibit the expressions of these genes, but interestingly, they were not reduced in *WT_CUR*. In addition (Fig6. M, N), MAOA and MAOB reduce molecular oxygen when oxidizing substrates and produce ROS, so their high expression may cause oxidative stress. Similarly, their expression was CUR-inhibited only in *Nrf1α*^*–/–*^_*CUR* tumor tissues.

## 4. Discussion

Relevant research indicates that Nrf1 serves to inhibit tumor progression and assumes a pivotal role in growth and development. Under normal physiological conditions, Nrf2 functions to safeguard cells against DNA damage induced by ROS and carcinogens, thereby impeding tumorigenesis. However, within cells bearing carcinogenic mutations, Nrf2 aids cells in withstanding stress, consequently sustaining the survival of cancer cells. The MTT results show that CUR has a more potent inhibitory effect on the growth of WT cells. This suggests that Nrf1/2 might be crucial for the function of CUR. Regarding the subsequent intervention time and concentration of CUR, in both WT and *Nrf2*^*–/–*^ cells, CUR promoted the expression of Nrf1 protein. In WT cells, the expression of Nrf2 protein initially increased and then decreased (which is consistent with the Nrf2 *Luc* results, Fig S1. C). However, in *Nrf1α*^*–/–*^ cells, the expression of Nrf2 gradually declined. These findings indicate that CUR exhibits a concentration and time - dependent trend on the protein levels of Nrf1 and Nrf2. Moreover, CUR follows a bidirectional regulatory pattern on Nrf2 expression, suggesting that CUR has a balancing effect on excessive stress. Simultaneously, we discovered that CUR exerted a potent activating effect on the transcription and expression of the downstream genes of Nrf1/2, namely HO - 1, GCLC, GCLM, GSR, and GPX1. However, *Nrf2*^*–/–*^ cells were scarcely influenced by CUR. At the protein level, CUR promoted the expression of the aforementioned genes in WT cells. In contrast, this promotional effect was attenuated or not significant in *Nrf1α*^*–/–*^ cells, and the changes in *Nrf2*^*–/–*^ cells were rather modest. These results demonstrate that CUR can regulate the mRNA and protein expression of downstream target genes via Nrf1/2. It also mitigates the aberrant expression of these genes in *Nrf1α*^*–/–*^ cells. Moreover, Nrf2 deficiency, to a certain extent, retards the effectiveness of CUR.

ROS, serving as a crucial signaling molecule, are implicated in diverse cellular life activities, including cell growth, proliferation, differentiation, survival, DNA damage induction, and apoptosis. As a result, compared to Nrf1 or Nrf2 knockout cells, CUR more effectively promoted DNA damage, apoptosis, and ROS generation in WT cells. However, CUR did not elicit significant alterations in the aforementioned phenotypes within *Nrf2*^*–/–*^ cells. These results imply that CUR likely induces DNA damage and apoptosis in hepatocellular carcinoma cells by promoting ROS production. This phenomenon is also observed in *Nrf1α*^*–/–*^ cells, yet not in *Nrf2*^*–/–*^ cells. These findings indicate that the apoptosis and damage of cancer cells induced by CUR are predominantly mediated by Nrf2. Regarding the antioxidant aspect, the results of CUR intervention demonstrated that the absence of Nrf1 and Nrf2 attenuated the effect of CUR to a certain degree. In fact, in *Nrf1α*^*–/–*^ cells, CUR even downregulated the protein levels of overexpressed genes, including G6PD, HO-1, TXN2, and SOD2. CUR is regarded as both an antioxidant and a hormetin. Its regulation of the stress response adheres to a bidirectional pattern ^[22]^. On one hand, it can counteract the generation of ROS and safeguard cells from oxidative stress disturbing their normal state. On the other hand, it has the ability to enhance the oxidative stress in cancer cells and trigger their entry into the apoptosis process ^[23]^.

In the events following ROS accumulation, GRP78, a target gene in the UPR process, can bind to three ER sensor proteins, namely IRE1, PERK, and ATF6, rendering them inactive. Once dissociated from GRP78, IRE1 and PERK form homodimers respectively and initiate downstream signaling cascades via phosphorylation ^[24]^. CUR is capable of activating the mRNA and protein expression of GRP78 and its downstream target genes in cells. This indicates that CUR can augment ER stress and the UPR response in both WT and *Nrf1α*^*–/–*^ cells. Additionally, it has been reported that protein toxicity and oxidative stress resulting from elevated ROS levels can cause the accumulation of misfolded or unfolded proteins in the mitochondrial matrix, thereby activating the UPR^mt [25]^. In the presence of CUR, eIF2α and ATF4 also mediated the expression of CHOP in mitochondria, thereby promoting the occurrence of the UPR. Intriguingly, in *Nrf2*^*–/–*^ cells, the expression of Fibroblast growth factor (FGF21) was significantly elevated. FGF21, which is abundant in the human liver, is primarily involved in the regulation of glucose, lipid, and amino acid metabolism. It can upregulate its own expression in response to various stressors. When liver cancer or irreversible liver diseases occur, FGF21 can be highly induced to protect the liver from damage; conversely, its lack of expression can lead to inflammation ^[26]^. In this study, the basal level of FGF21 was extremely low in both WT and *Nrf1α*^*–/–*^ cells, with the level being particularly low in *Nrf1α*^*–/–*^ cells. In contrast, the basal expression of FGF21 was notably high in *Nrf2*^*–/–*^ cells, and its expression was significantly up - regulated following CUR intervention. These findings seemingly suggest the role of Nrf1 in inflammation and tumor progression, as well as the potential of CUR in counteracting inflammation.

Simultaneously, CUR is still capable of promoting the expression of genes associated with tumor signaling pathways. Relevant studies suggest that CUR can promote cancer cells to enter the death process through ROS generation and stress response, and exogenous and endogenous death pathways are not excluded in different cell lines. Firstly, CUR significantly promoted the proteins expression and phosphorylation of JNK, ERK1/2, P38 and others, and also up-regulated the expression of c-JUN, which also occurred in WT and *Nrf1α*^*–/–*^ cells; however, the phosphorylation level of MEK and c-JUN expression were significantly decreased in *Nrf2*^*–/–*^ cells. In addition, CUR also induced the up-regulation of AKT and PI3K, but also promoted the expression of PTEN in WT and *Nrf2*^*–/–*^ cells. These phenomena indicate that CUR can also affect the expression of key genes in AMPK and PI3K/AKT signaling pathway to alleviate tumor through regulation of oxidative stress-related genes, and Nrf1/2 is also required to mediate this process. Furthermore, CUR could remarkably reduce the migration of liver cancer cells in both WT and *Nrf1α*^*–/–*^ cells. Notably, the expression of migration-related genes exhibited more pronounced changes in *Nrf1α*^*–/–*^ cells. This suggests that CUR requires Nrf2 to mediate the inhibition of tumor cell migration. Although the specific regulatory mechanism between cell migration and Nrf1 remains incompletely understood, recent research has reported the correlation between Nrf2 and autophagy ^[26]^. Meanwhile, members of our research group are continuously delving into the regulatory relationship between autophagy in liver cancer and Nrf1/2.

CUR relies on Nrf2 to mediate various molecular mechanisms or phenotypes that are otherwise associated with Nrf1. Although CUR’s activation of Nrf1 persists, this might be linked to the concurrent activation of Nrf2. Interestingly, CUR is still capable of activating the expression of Nrf1 in *Nrf2*^*–/–*^cells. These findings suggest that CUR may induce Nrf1 expression via molecular pathways distinct from those involving Nrf2. Furthermore, the transcriptome data (Fig S2. A-F) reveal that CUR can still exert its effects via the *ALPI, MCM7, TXNIP, ALPP* and *SMIM11A* genes (notably *ALPP* and *MCM7*), irrespective of the absence of Nrf1 or Nrf2. In the absence of Nrf1, CUR predominantly regulates cell fate through the hepatitis B, microRNA, and tumor signaling pathways. Among these, Nrf1 can mediate CUR - induced regulation of the expression of the *UTP14C, FLRT1* and *PNPLA1* genes. Similarly, as shown in Fig S2. J-L, Nrf2 can mediate the expression of the *DUSP8, EFNA1, HSPA1A, HSPA1B, HSPA8* and *TGFβ2* genes within the MAPK signaling pathway regulated by CUR.

In conclusion, in WT cells, CUR up-regulated the expression of antioxidant genes by activating Nrf1/2 and their downstream target genes. Concurrently, it promoted ER stress (ROS accumulation), enhanced the expression of PTEN and p - p38, and down - regulated the MMP9 and SNAIL1. These actions alleviated tumor development through the activation of PERK and ATF6, ultimately promoting cancer cell apoptosis and impeding their growth. In *Nrf1α*^*–/–*^ cells, the inhibition of Nrf2 expression affected the expression of downstream target genes. The down - regulation of ATF6 and eIF2α relieved ER stress, while mitochondrial stress (ROS aggregation) was up - regulated by CHOP and FGF21. Additionally, the inhibition of PI3K, promotion of p - p38, down - regulation of SNAIL1 and ITGB4 alleviated tumor growth. Finally, in combination with DNA damage, apoptosis was promoted. Conversely, in *Nrf2*^*–/–*^ cells, CUR’s regulation of different genes was generally not significantly different. Overall, the differential responses of HepG2 cells with different genotypes to CUR were mainly ascribed to the absence of Nrf1.

In a recent study, Qiu et al. also discovered that the loss of Nrf1 in human HepG2 cells can result in cell death following glucose deprivation. They posited that the glycosylation of Nrf1 endows it with the ability to sense the energy state, thereby confirming the role of Nrf1 as a dual receptor and regulator during glucose homeostasis ^[27]^. Moreover, in *Nrf1α*^*–/–*^ HepG2 cells, the basic expression of genes in the AMPK signaling pathway was inhibited. The same findings were obtained in our study, suggesting a potential metabolic equilibrium between Nrf1 and glucose metabolism as well as energy metabolism. The results presented that, in the context of Nrf1/2 - mediated oxidative stress damage and tumor signaling pathways, CUR exerts its effects mainly through Nrf2. However, the impact of CUR on the metabolic pathways within this system remains unclear. In cells (Fig S3. A), we initially observed that CUR significantly diminished the formation of lipid droplets in the three genotypes of HepG2 cells, with a particularly notable difference in *Nrf1α*^*–/–*^ cells. Consistent results were also evident in transplanted tumor tissues (Fig S3. B), where the formation of lipid droplets in the xenografted tumor tissues of *Nrf1α*^*–/–*^ cells was significantly reduced. However, in WT cells, CUR had no significant impact on processes such as glucose level, ATP ratio and NADPH transformation. In contrast, in the tumor tissues of *Nrf1α*^*–/–*^ cells, these parameters decreased in significantly. This indicates that CUR is more likely to target lipid synthesis and metabolism within cells. Meanwhile, in xenografted tumors, CUR exerts significant effects on energy and nutrient metabolism. It is notable that both TRP metabolism (Fig6. G) and glutathione metabolism in which CUR is involved (Fig S4. E - I) in tissues are relevant metabolic pathways. When Nrf1 is deficient, these pathways can induce the malignant development of tumors. Here, CUR more markedly regulated the expression of key metabolite enzymes. In particular, it inhibited the mRNA levels of *MAOB, KYNU* and *IDO2* enzymes. As a result, it reduced the accumulation of the metabolite Kyn and regulated glutathione synthesis. However, the inhibitory effect of CUR on the growth of xenografted tumors *in vivo* was not significant. Regardless of the presence or absence of Nrf1, the multiple reactions or signaling occurring in xenografted tumor tissues were not significantly manifested in the tumor itself.

According to reports, when CUR is taken orally, the majority of it is excreted in the feces. Only a small fraction is absorbed in the intestine and is then rapidly metabolized in the plasma and liver. The poor oral bioavailability of CUR can be attributed to three physiological barriers in the gastrointestinal tract. Firstly, there is a physical barrier that restricts drug transport and impedes the spread of CUR ^[28]^. Secondly, gastric acid, bile, and various digestive enzymes can lead to the degradation of CUR in bile, thereby affecting its absorption into the bloodstream (chemical barrier) ^[29]^. Thirdly, metabolic enzymes and epithelial P-glycoprotein (P-gp) efflux can inactivate CUR and limit its gastrointestinal absorption (biochemical barrier) ^[30]^. Therefore, we surmise that the above - mentioned results might be associated with multiple factors, including the physical and chemical properties of CUR, the mode of administration, and the dosage. In addition, CUR did not induce Nrf1 expression in WT tumor tissues, but still significantly inhibited Nrf2 overexpression in tumors with Nrf1 deletion, indicating that CUR affects the regulation of KYNU, MAOB and other metabolism-related genes in xenograft tumor tissues mainly through Nrf2 mediation. The excessive accumulation of kyn and GSH can be alleviated. However, this phenomenon implies that CUR, as a homeostasis stabilizer, has an unexpected negative regulatory effect on Nrf2 that affects redox homeostasis, because related studies have suggested that CUR is an Nrf2 activator^[31-33]^ (in both normal and tumor/inflammatory cells). However, in the present study, overexpressed Nrf2 was still suppressed by CUR after Nrf1 knockout (at least in the *Nrf1α*^*–/–*^ cells and xenografted tumor).

## 5. Conclusion

Herein, both Nrf1 and Nrf2 mediated the effects of CUR on the phenotypes of different genotypes of cells. *Nrf1α*^*–/–*^ cell demand for energy (ATP) and nutrition (glucose) and related genes (*SREBP, FASN, HK1*, etc.) were significantly down-regulated by CUR under Nrf2, thus regulating metabolic progress. In xenograft tumors, CUR can mitigate the abnormal metabolism of tryptophan or glutathione in *Nrf1α*^*–/–*^ tumor tissues. Additionally, CUR acts through Nrf1/2 to differentially regulate the expression of proteasome subunits and uphold the stability of the Nrf1 protein. As a result, CUR not only regulates proteasome homeostasis but also influences Nrf1’s role in coordinating redox homeostasis. In recent years, although there have been many studies that CUR is a targeted activator of Nrf2 and participates in the treatment and defense of a variety of diseases, CUR is not a Nrf2 activator in a strict sense in this study. The specific manifestations are as follows: overexpression of Nrf2 in *Nrf1α*^*–/–*^ cells can be significantly inhibited by CUR in a concentration or time-dependent manner, and this phenomenon has also been described in previous studies related to drug intervention^[17]^, which provides new ideas for the regulatory relationship between CUR and Nrf2. Simultaneously, CUR exhibits a definite activation effect on Nrf1, yet lacks precise targeting. In this regard, we consider that employing chemical methods to optimize the structure of CUR and enhance its targeting specificity is crucial. Thus, the relevant work is also being actively carried out.

## Author contributions

R.W. designed and performed most of the experiments, collected all the relevant data, and wrote a draft of this manuscript with most figures and supplemental information. J.F. performed the subcutaneous xenograft tumor experiments and some of drug intervention. Both S.H. and M.W. helped R.W. with mice anatomy, tissue section, metabolomics and transcriptome data analysis and other experiments. And L.K. helped R.W performed some of the western blotting and data collection. Lastly, Y.Z. designed and supervised this study, analyzed all the data, helped to prepare all figures with cartoons, rewrote and revised the paper. All these co-authors have read and agreed to the published version of the manuscript.

## Supplementary Materials

The supporting information, including four supplemental figures and also one supplemental table.

## Acknowledgments

We are greatly thankful to Dr. Yonggang Ren (North Sichuan Medical College, Sichuan, China) and Dr. Lu Qiu (Zhengzhou University, Henan, China) for their involvement in establishing the indicated cell lines used in this study, and also thank to all other present and past members of Zhang’s laboratory for giving critical discussion and invaluable help with this work. ***Foundation:*** This study was funded by the National Natural Science Foundation of China (NSFC project 82473147, 82073079) awarded to Prof. Yiguo Zhang (at Chongqing University). This is also, at part, supported by the Natural Science Foundation (2024D01C58) of Xinjiang Uygur Autonomous Region awarded to Prof. Reziyamu Wufuer (at Xinjiang university).

