## supplement materials (Fig S1 to S4) for "Curcumin activates distinct programming of redox metabolism towards differential regulation of gene expression mediated by Nrf1 and Nrf2"

### Curcumin affects Nrf1/2-mediated redox-metabolism-differential regulation of gene expression by activating Nrf1 (Nfe2l1)

**Reziyamu Wufuer[1,2#], Feng Jing[1,3#], Hu Shaofan[1,3], Wang Meng[1,4], Liu Keli[1], Chen Xi[1], Zhang Yiguo[1,5*]**

[1]The Laboratory of Cell Biochemistry and Topogenetic Regulation, College of Bioengineering and Faculty of Medical Sciences, Chongqing University, No. 174 Shazheng Street, Shapingba District, Chongqing 400044, China

[2]School of Parmaceutical Sciences and Institute of Materia Medica, Xinjiang University, Xinjiang, 830017, China

[3]Jinfeng Laboratory, No. 313 Jinyue Road, Chongqing High-tech District, 401329, China

[4]Department of Pathophysiology, College of High Altitude Military Medicine, Third Military Medical University (Army Medical University), No. 30 Gaotanyan Street, Shapingba, Chongqing, 400038, China

[5]School of Life and Health Sciences, Fuyao University of Science and Technology, No. 104 Wisdom Avenue, Nanyu Town, Minhou County High-Tech District, Fuzhou, 350109, Fujian, China

[#] Those authors equally contributed to this paper.

#### Supplement materials:

**Figure. S1**

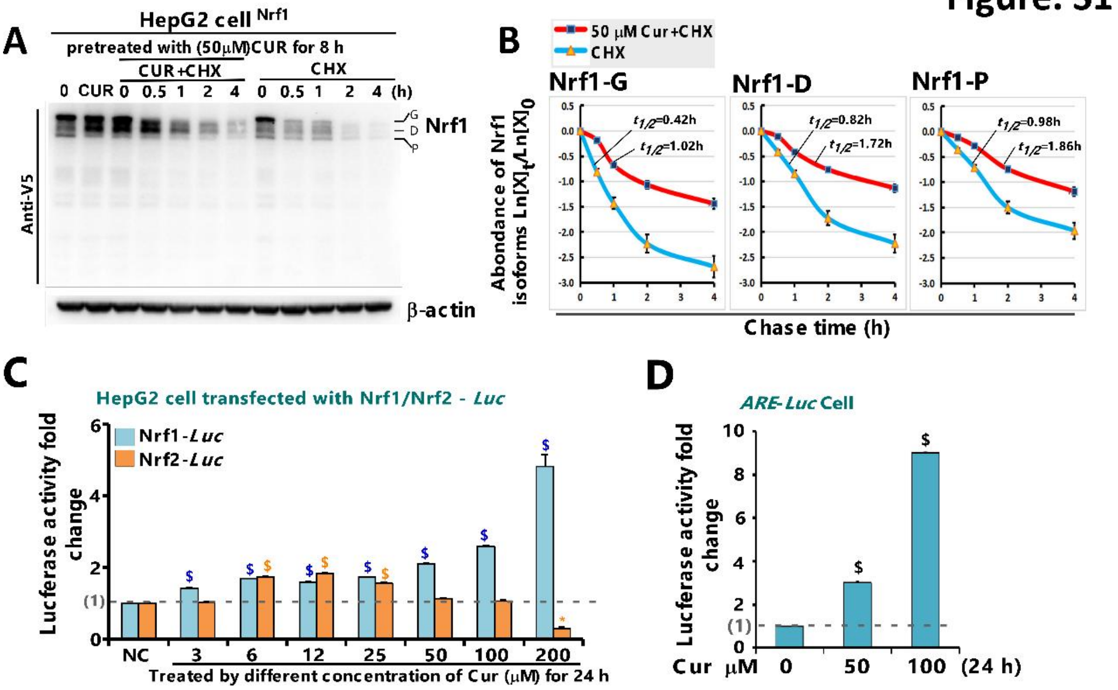


**Figure S1.** ***CUR promotes Nrf1/2 transcriptional activity and exogenous Nrf1 protein stability.*** **(A)** The changing of the stability of Nrf1α protein after 50 μM CUR stimulation, **(B)** and the expression level of each protein band is represented by a time curve . All protein expression above were determined by western blotting with the indicated antibodies. In addition, the expression of exogenous Nrf1α after 50 μM CUR stimulation is realized by transferring Nrf1 plasmid (empty as the control group) in HepG2 (*WT*) cells. The intensity of relevant immunoblots representing different protein expression levels was also quantified by the Quan-tity One 4.5.2 software. The resulting data were then shown graphically, after being calculated by a formula of *Ln ([X]t/[X]0)*, in which $[X]_t$ indicated a fold change (mean±SD) in each of those examined protein expression levels at different times relative to the corresponding controls measured at 0 h (i.e., $[A]_0$), relative to their (CUR+ CHX) or (CHX) control, respectively, and which were representative of at least three independent experiments. **(C)** The transcriptional activity of Nrf1/2 luciferase reporter gene by CUR at different concentrations (0-200 μM) was detected in *WT* cells, and the *Nrf1-Luc* plasmid was constructed in the laboratory, through which we could observe the transcriptional activity of *Nrf1* by CUR. **(D)** The transcriptional activity of *Nrf1-Luc* could be enhanced by CUR intervention in the range of 3-200 μM, and the transcriptional activity of *Nrf2-Luc* was enhanced at the concentration of 6-25 μM, and decreased at the concentration of 200 μM. These results indicated that CUR strongly promoted the transcription of Nrf1 luciferase reporter gene in *WT* cells, and the results were consistent with those of *ARE-Luc* stable expression cells (*ARE-Luc* cell line was constructed by literature method and is stored in our laboratory (DOI: 10.1158/0008-5472.CAN-06-2298)).

**Figure S2**

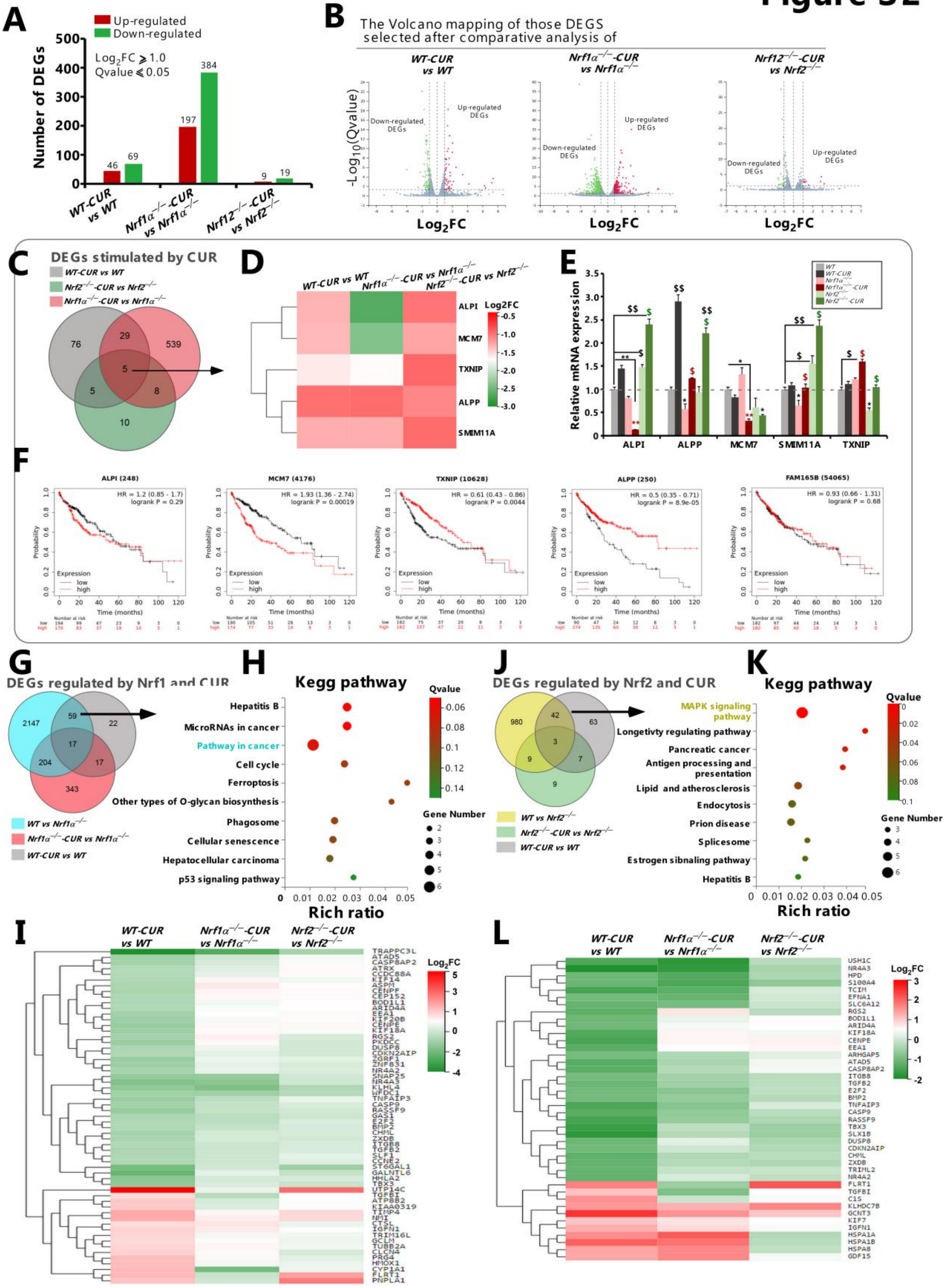


**Figure S2.** ***The pharmacological effect of CUR on different genotype of HepG2 cells is mainly achieved by stimulating target genes regulated by Nrf2.*** **(A)** Distinct regulation of those differential expression genes by CUR in WT, *Nrf1α*$^{-/-}$ and *Nrf2*$^{-/-}$ cell lines were further analyzed. The intervened effects of CUR on such genes were statistically determined relative to their corresponding vehicle controls, and then shown graphically. **(B)** Those differentially regulated genes were illustrated by distinct volcano plots. Red represents gene up-regulation by CUR, green denotes gene down-regulation by CUR, whereas gray shows no-significant differences. **(C)** The Venn plot shows distinct cell group-intersected genes stimulated by CUR, which were selected mainly by calculating the parameter of CUR-induced fold changes (i.e., $| Log_2FC | \geq 1.0$). **(D)** The Venn plot shows those genes mediated by Nrf1 and Nrf2 after CUR intervention, (E) and their differential expression changes were also further presented within distinct heat-maps, after being comparatively analyzed in the three pairs of differently treated cell lines. **(F)** Real-time qPCR analysis of differential gene expression stimulated by CUR. The resulting data were shown as fold changes (mean±SD, n = 3×3), which are representative of at least three independent experiments being each performed in triplicates. Among them, significant increases ($, $p < 0.05$;$$, $p < 0.01$) and significant decreases (*$p < 0.05$), were statistically determined, when compared with their corresponding *WT* controls. Additional symbols "$ or *" indicate significant differences in *Nrf1α*$^{-/-}$ or *Nrf2*$^{-/-}$ cell lines, relative to their $T_0$ values after CUR intervention. **(F)** And, the impact of these deferentially expressed genes on the prognosis survival percentage of liver cancer patients. **(G-I)** After KEGG enrichment of 59 genes from Venn plots, it was found that more genes were enriched in the pathway in cancer **(G,H)**. The related genes were *E2F2* (E2F Transcription Factor 2, E2F Transcription Factor 2), *HO-1* (HMOX1), *BMP2* (Bone Morphogenetic Protein 2, a ligand that can bind to *TGFβ* family proteins), *TGFβ2* (Transforming Growth Factor Beta 2), *Caspase9* (Cysteine Aspartate Protease Family Protein 9, involved in cell apoptosis pathway), and *CCNE2* (Cycle E2, involved in cell cycle regulation), which are respectively involved in cell proliferation, apoptosis, and cell cycle regulation. In contrast, the three genes *UTP14C* (involved in the cell cycle), *FLRT1* (leucine rich fibronectin transmembrane protein), and *PNPLA1* (key lipid metabolism gene) were all elevated in *WT* and *Nrf2*$^{-/-}$ cells, while there was no difference in *Nrf1α*$^{-/-}$ cells, indicating that they seem to be more closely related to *Nrf1*. **(J-L)** Similarly, Venn diagram intersection analysis was conducted on the target genes or signaling pathways regulated by CUR through *Nrf2*. The results showed that a total of 42 genes could be regulated by CUR **(J,K)**. The heatmap results showed that CUR had more differential gene regulation in *WT* and *Nrf1α*$^{-/-}$ cells **(L)**. More genes in the top ten pathways are enriched in the MAPK signaling pathway, including *DUSP8* (Dual Specification Phosphatase 8), *EFNA1* (Ephrin A1, EPH family protein), *HSPA1A* (Heat Shock Protein Family A Member 1A, Heat Shock Protein 70 subfamily protein), *HSPA1B* (Heat Shock Protein Family A Member 1B, Heat Shock Protein 70 subfamily protein), *HSPA8* (Heat Shock Protein Family A Member 8, Heat Shock Protein 70 subfamily protein), *TGFβ2* (Transforming Growth Factor Beta 2, Transforming Growth Factor β2), etc. Participate in cell proliferation/apoptosis or protein synthesis processing. Transcriptome data showed that *DUSP8* and *TGFβ2* were significantly down regulated in *WT* cells after CUR intervention, while other genes were significantly up-regulated in *WT* and *Nrf1α*$^{-/-}$ cells, with no differential changes observed in *Nrf2*$^{-/-}$ cells. Taken together, these results demonstrate that *Nrf1* and *Nrf2* are differentially contributable to distinct effects of CUR on those genes,

### Figure.S3

Oil red staining of different type of HepG2 cells and xenograft tumor tissues

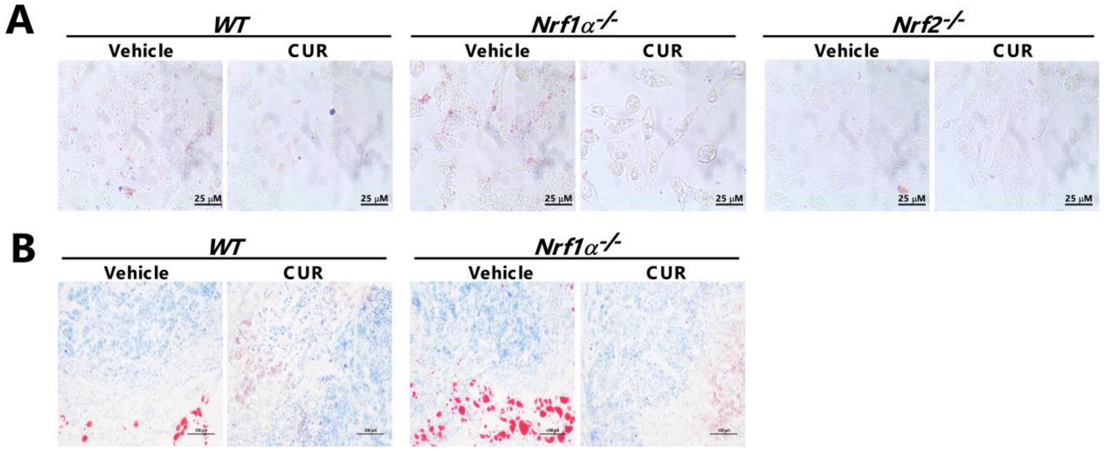


**Figure S3.** ***CUR can inhibit lipid droplet deposition in liver cancer cells and xenograft tumor tissues**.* **(A)** First, treat cells of different genotypes with CUR for 0 or 24 h, discard the supernatant, stain with Oil Red O for 20 min, wash with PBS until the background is colorless, and finally observe and record under a regular microscope. **(B)** The pre-made sections of tumor tissues were stored in a -20℃ refrigerator on a slicing rack. Before experiment, they were allowed to warm up for 5-10 min in room temperature. Thereafter, oil red O staining of these tumor sections was carried out according to the manufacturer's instructions and then subjected to microscopic observation of red lipid droplets (D027-1, Nanjing Jiancheng, Nanjing, China).

### Figure.S4

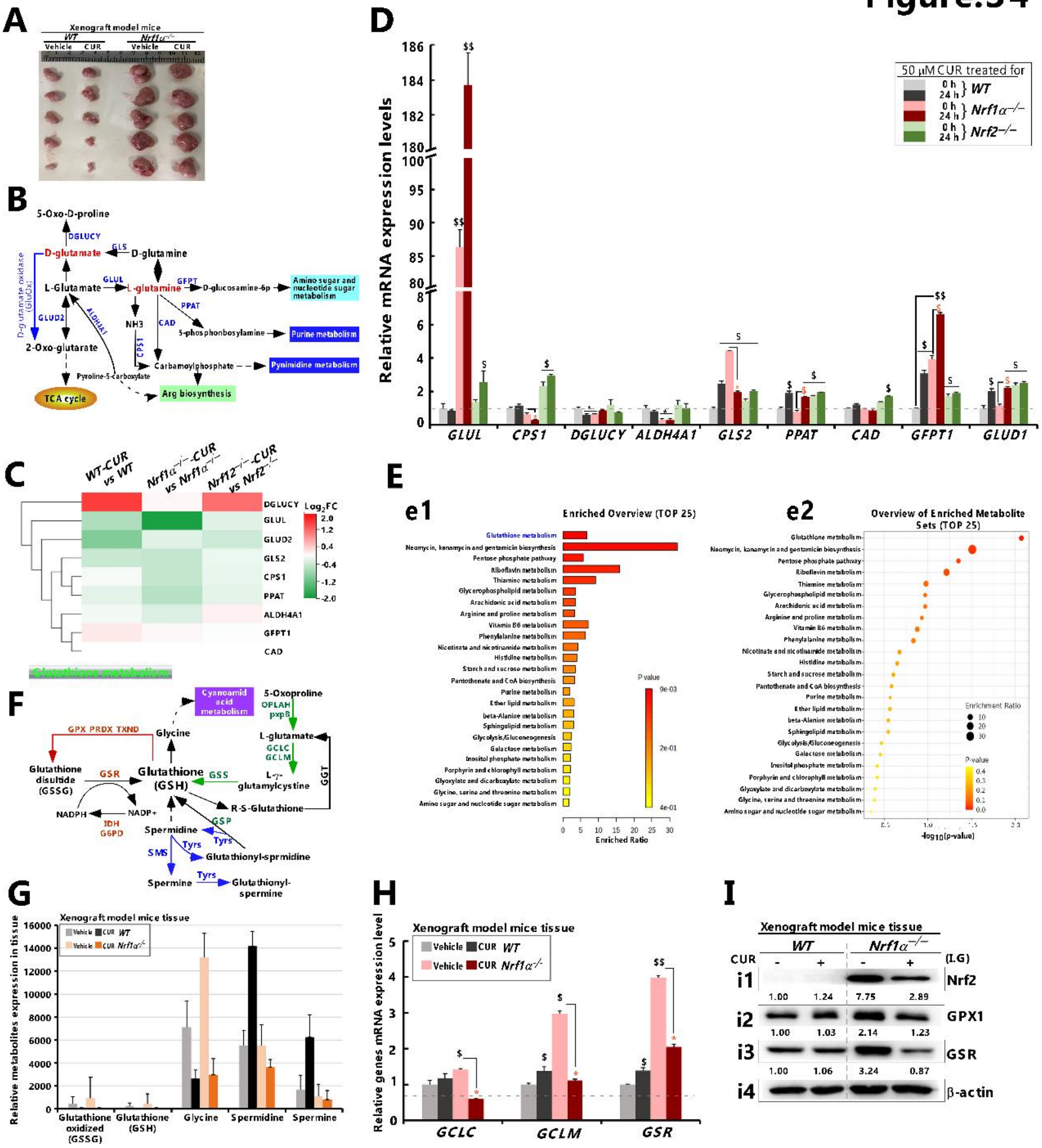

**Figure S4.** ***CUR affected xenograft tumor weakly but still affected D-glutamate and glutathione metabolic process.*** **(A)** The malgrowth phenomenon of xenograft tumors tissues in distinct groups of nude mice, that were subcutaneously injected with human *WT* or *Nrf1α*$^{-/-}$ hepatoma cell lines, followed by intervention with CUR (I.G., 35 mg/mL) or vehicle (I.G., 0.9% physiological saline). **(B)** After conducting untargeted metabolomics analysis on cells of different genotypes, we found that there were 879 compounds with completely opposite expression trends when Nrf1 and Nrf2 were present (i.e., | $Log_2FC$ | ≥1.5). After enrichment analysis using the HMDB database, significant differences were observed in the D-glutamate metabolic pathway, and metabolic pathways were plotted based on the KEGG database, where "red" represents the key differential metabolites in the pathway and "blue" represents the key enzymes in the metabolic pathway. **(C)** Subsequently, gene expression heat maps and **(D)** mRNA detection results of related metabolic enzymes showed significant differences in the basal levels of genes such as *GLUL, CPS1, GFPT, GLUD1* in cells of different genotypes, and there was a significant trend of change in cells treated with CUR. **(E)** In addition, we also used a similar method to enrich metabolites in tumor tissues and cells (note: the selection criteria for differential metabolites are "only Nrf1 exists and there is differential expression after CUR intervention", | $Log_2FC$ | ≥1.5). **(F)** The results showed that the glutathione metabolism pathway had the most significant differences, and the KEGG database was used to plot the direction of this metabolic pathway. **(G)** GSSG, GSH, Glycine has higher basal levels in *Nrf1α*$^{-/-}$ tissues, and there is no difference between spermidine and arginine in *WT*, but after CUR intervention, GSSG, GSH, Glycine were significantly reduced (* $p$<0.05). **(H-I)** In addition, the mRNA and protein levels of enzymes GCLC, GCLM, GSR, and GPX1 related to GSH synthesis were much higher in *Nrf1α*$^{-/-}$ tissues than in *WT*. Although the expression of GCLC protein was not significant in *WT* after CUR treated, the mRNA and protein expression of all genes were significantly reduced in *Nrf1α*$^{-/-}$ tumor tissue (* $p$<0.05), and Nrf2 was much higher in *Nrf1α*$^{-/-}$ tissues than in *WT* (note: HepG2 cells did not form tumors *in vivo* after Nrf2 knockout). Real-time qPCR analysis of differential gene expression stimulated by CUR. The resulting data were shown as fold changes (mean±SD, n = 3×3), which are representative of at least three independent experiments being each performed in triplicates. Among them, significant increases ($, $p$ < 0.05;$$, $p$ < 0.01) and significant decreases (*$p$ < 0.05), were statistically determined, when compared with their corresponding *WT* controls. Additional symbols "$ or *" indicate significant differences in *Nrf1α*$^{-/-}$ or *Nrf2*$^{-/-}$ cell lines, relative to their $T_0$ values after CUR intervention. The intensity of relevant immunoblots representing different protein expression levels was also quantified by the Quan-tity One 4.5.2 software.
