## supplemental table 1. for "Curcumin activates distinct programming of redox metabolism towards differential regulation of gene expression mediated by Nrf1 and Nrf2"

### Curcumin affects Nrf1/2-mediated redox-metabolism-differential regulation of gene expression by activating Nrf1 (Nfe2l1)

**Reziyamu Wufuer[1,2#], Feng Jing[1,3#], Hu Shaofan[1,3], Wang Meng[1,4], Liu Keli[1], Chen Xi[1], Zhang Yiguo[1,5*]**

[1]The Laboratory of Cell Biochemistry and Topogenetic Regulation, College of Bioengineering and Faculty of Medical Sciences, Chongqing University, No. 174 Shazheng Street, Shapingba District, Chongqing 400044, China

[2]School of Parmaceutical Sciences and Institute of Materia Medica, Xinjiang University, Xinjiang, 830017, China

[3]Jinfeng Laboratory, No. 313 Jinyue Road, Chongqing High-tech District, 401329, China

[4]Department of Pathophysiology, College of High Altitude Military Medicine, Third Military Medical University (Army Medical University), No. 30 Gaotanyan Street, Shapingba, Chongqing, 400038, China

[5]School of Life and Health Sciences, Fuyao University of Science and Technology, No. 104 Wisdom Avenue, Nanyu Town, Minhou County High-Tech District, Fuzhou, 350109, Fujian, China

[#] Those authors equally contributed to this paper.

#### Table 1. The primer pairs used for qRT-PCR analysis.

| Gene ID | Accession number | Name | Forward primers(5'-3') | Reverse Primers (5'-3') |
|---|---|---|---|---|
| 60 | NM_001101 | *β*-actin | CATGTACGTTGCTATCCAGGC | CTCCTTAATGTCACGCACGAT |
| 4779 | NM_001330261 | *Nrf1* | GAAGCCCACCAAGACCGAA | GCCTCTTCCTGTACACTGACC |
| 4780 | NM_001313903 | *Nrf2* | ATATTCCCGGTCACATCGAGA | ATGTCCTGTTGCATACCGTCT |
| 11047 | NM_007002.4 | ADRM1 | TACATTCAGCAGACGGACGAC | AACTTCAGCACGTAGACCCT |
| 5686 | NM_002790.4 | PSMA5 | CCATGTTTCTTACCCGGTCT | TTCTCCACAGCTAGGCACAC |
| 5689 | NM_002793.4 | PSMB1 | GCGATGTTGTCCTCTACAGC | TGCCAGTATAGTACCTCCGTTG |
| 5690 | NM_002794.5 | PSMB2 | TCTGCCAACCTTCAGTGTTCG | CCATTTATCAAGAGTGCGCCTG |
| 5691 | NM_002795.4 | PSMB3 | CACTGACGTCCAGACAGTTGC | ACCGTTTCTCATACAAGAGGTT |
| 5694 | NM_002798.3 | PSMB6 | CCTACCAGCTCGGTTTCCAC | TGATTCCCGCCATCAGGTCT |
| 5700 | NM_002802.3 | PSMC1 | CCCTGCCTGATGAAAAGACGAA | TTAGCCATGATCAGGTCGTCCA |
| 5692 | NM_153001.3 | PSMC4 | GCCACAAACAGAGCAGACACC | GGGCCACATAGTCTTCCAAGTCA |
| 5705 | NM_001199163.2 | PSMC5 | TCGAGCTGCCTGTTAAGCATC | TTCAGAGCCAGAGACACGAA |
| 5707 | NM_002807.4 | PSMD1 | GCGTCCTTGTATTTACCCAGT | AACTTTCGGCATCTTTAAGTCC |
| 2708 | NM_002808.5 | PSMD2 | CCTCCTCTTTTGTGAATGGCTT | TGCACTCAACATTCCGTGGT |
| 5709 | NM_002809.4 | PSMD3 | AGGAAACCTAGCCAAGTTCAACC | GGCTGATCATGCGTACACCT |
| 5710 | NM_001330692.2 | PSMD4 | ACCGACAAGGCAAGAATCACA | ACTTTCTCCTTCTTGAGGCGTTT |
| 9861 | NM_001271779.2 | PSMD6 | GCCTTTCGCAAGACATATGACA | TTGGCCTTTTCTGTGTTTCGT |
| 5719 | NM_002817.4 | PSMD13 | GAGATGACTTTCACACGACCT | TGTCCACCTCGTCTATACTGC |
| 3162 | NM_002133 | *HO-1* | CAGAGCCTGGAAGACACCCTAA | AAACCACCCCAACCCTGCTAT |
| 1728 | NM_001025434 | *NQO1* | AAGAAGAAAGGATGGGAGGTGG | GAACAGACTCGGCAGGATACTG |
| 2729 | NM_001197115 | *GCLC* | TCAATGGGAAGGAAGGTGTGTT | TTGTAGTCAGGATGGTTTGCGA |
| 2730 | NM_001308253 | *GCLM* | TGCTGTGTGATGCCACCAGA | CGCTTGAATGTCAGGAATGCTT |
| 2936 | NM_001195102 | *GSR* | CACGAGTGATCCCAAGCCC | CAATGTAACCTGCACCAACAATG |
| 2876 | NM_001329503 | *Gpx1* | CAGTCGGTGTATGCCTTCTCG | GAGGGACGCCACATTCTCG |
| 5034 | NM_000918 | *P4HB* | ATCTTCATCGACAGCGACCACACCG | CGGTGTGGTCGCTGTCGATGAAGAT |
| 3309 | NM_005347 | BIP | AAAGCCACCAAGATGCTG | GCCTGCACTTCCATAGAGT |
| 9451 | NM_001313915 | *PERK* | TGCTTCTACAGCGTACCCAA | TCAATAAATCCGGCTCTCGT |
| 83939 | NM_001319045.2 | EIF2A | ACAATCAGGAAACGATAAGCCAT | TGCCAAATCTGGACTCTTGTCAC |
| 468 | NM_182810 | ATF4 | CCCTTCACCTTCTTACAACCTC | TGCCCAGCTCTAAACTAAAGGA |
| 2081 | NM_001433 | *IRE1* | AGTCTCTGCCCATCAACCTC | GCATTCCACCGGAGCTCTCG |
| 7494 | NM_001394000 | *XBP1* | CACCCCTCCAGAACATCTCC | TGTCCAGAATGCCCAACAGG |
| 22926 | NM_007348 | ATF6 | CCCTCTCAGAAAACGAGCAAC | AGACAACTCTTCGCTTTGGAC |

| | | | | |
|---|---|---|---|---|
| 1649 | NM_004083 | *CHOP* | ACCAGCAGAGGTCACAAGCA | ATGACCACTCTGTTTCCGTT |
| 3329 | NM_002156.5 | HSP60 | GCCCTCCTTCGATGCATTCCA | ACCTGCATTCTTAGCAATGGTCA |
| 26291 | NM_019113.4 | FGF21 | GGAGCTTCTGCATCTATCCCAA | GTGCCAGATTCCAGTTGTCCA |
| 2539 | NM_000402.4 | G6PD | CGAGGCCGTCACCAAGAAC | GTAGTGGTCGATGCGGTAGA |
| 3417 | NM_005896.4 | IDH1 | ACACCAAGTGACGGAACCCAA | AGGCCAACCCTTAGACAGAGC |
| 3418 | NM_002168.4 | IDH2 | AGTGCGTTTTACCTCAGCCAGT | GTCCCACTACCTCCTCCCCTC |
| 4199 | NM_002395.6 | ME1 | GGGTCGGCTTTATCCTCCTT | ATGCTTCTTTGTTTTGCGGTT |
| 57103 | NM_020375.3 | TIGAR | TCACTGCCCTAAAAGTCCTCT | AAACTTCTGCCCATTCCCTC |
| 27244 | NM_014454.3 | SESN1 | CCTCAATGCTTAGACGGGCAA | TTCAGGAGTGCAAACAACAGT |
| 83667 | NM_031459.5 | SESN2 | CACCCAGACATGCTGTGCTT | TAGCCATGGTCTTCCCAGGT |
| 847 | NM_001752.4 | CAT | TTCTGTTGAAGATGCGGCG | GATGTAAAAAGTCCAGGAGGGGT |
| 5052 | NM_002574.4 | PRDX1 | TTCACTGACAAACATGGGGAA | CGCTCACTTCTGCTTGGAGA |
| 10935 | NM_006793.5 | PRDX3 | ATCTTGCCTGGATAAATACACC | TGATCTTAGTGCAAGACCAGA |
| 6647 | NM_000454.5 | SOD1 | TTGCATCATTGGCCGCACACT | ATTACACCACAAGCCAAACGACT |
| 6648 | NM_000636.4 | SOD2 | AATAGCTGGGATTACAGGTGCAT | CCAGACGGATCACTTGAGGTCAG |
| 2744 | NM_014905.5 | GLS1 | TTGGTCTTCCTGCAAAATCTGG | TCCCTTAACACTGTTGCCCAT |
| 23657 | NM_014331.4 | SLCA711 | TAGACATGTTCCATTCGAGGT | TGTACACAACTTTGCTGGTC |
| 7295 | NM_003329.4 | TXN1 | CCTTGCAAAATGATCAAGCCTT | TGGTGGCTTCAAGCTTTTCCT |
| 25828 | NM_012473.4 | TXN2 | TGACCACACAGACCTCGCCAT | ATGCCCACAAACTTGTCCACCAC |
| 140809 | NM_080725.3 | SRXN1 | GCTGTGTAACAAACCATCCCCAA | CAGCCCAAATTGCCAACCCAT |
| 207 | NM_001382430.1 | AKT1 | ACGGGCACATTAAGATCACAG | TCATTGTCCTCCAGCACCTCG |
| 2353 | NM_005252.4 | FOS | GACCGAGCCCTTTGATGACT | TCTGCTGCATAGAAGGACCCA |
| 4790 | NM_003998.4 | NFKB1 | GGGTAACTCTGTTTTGCACCT | AAACATGGCAGGCTATTGCT |
| 4791 | NM_001077494.3 | NFKB2 | CTTCTCTGCCTTCCTTAGAGCC | CCTGCTGTCTTGTCCATTCGAG |
| 5291 | NM_006219.3 | PI3K | ACTTACCAAGAATGGCTCGAT | CGTATTTACCCACGCTACAGG |
| 5728 | NM_000314.8 | PTEN | TAAAGATGGCACTTTCCCGTT | TTAGCCACTTCAGTTGGTGAC |
| 7157 | NM_000546 | TP53 | CCCACTTCACCGTACTAACCA | CATTAACCCTCACAATGCACT |
| 999 | NM_004360.5 | CDH1 | TTTTCCCTCGACACCCGAT | CCCACTGTATTCAGCGTGAC |
| 1000 | NM_001792.5 | CDH2 | AGAGTTTACTGCCATGACGTT | ACTGATTCTGTACACTGCGTTC |
| 6615 | NM_005985.4 | SNAI1 | CGTCCTTCTCCTCTACTTCAGTC | CTTTCGAGCCTGGAGATCCTT |
| 31 | NM_198834.3 | ACACA | GTTATTTCTCAGAGCTTCCGAAC | GAATTTCTTCTGCCAGTCCGAT |
| 1374 | NM_001876.4 | CPT1a | TGCAAAGGCGACATCAATCCG | AAATCCACGTCGTTTGCCAGA |
| 2194 | NM_004104.5 | FASN | GCCTGCCACAACTCCAAGGAC | TTCCTCAGCTGCTCCACGAAC |
| 6720 | NM_004176.5 | SREBP1 | GGAGCCATGGATTGCACTTT | CAGGAAGGCTTCAAGAGAGG |
| 6721 | NM_004599.4 | SREBP2 | CGACTCTGACCAGCACCCACA | GACGCTCAGGACAATCACACC |
| 6319 | NM_005063.5 | SCD | CACCACATTCTTCATTGATTGC | TCAGCCACTCTTGTAGTTTCCA |
| 6541 | NM_003045.5 | SLC7A1 | TCCCCATCCTGAGCATCTTC | CAGCATCCACACAGCAAACC |
| 6513 | NM_006516.4 | GLUT1 | TCCAGCTGCCATTGCCGTTG | AGGGACCACACAGTTGCTCCAC |
| 6514 | NM_000340.2 | GLUT4 | TGACCAGATCTCAGCTGCCTT | CCTGGCCCCTCAGTCGTTC |
| 3098 | NM_033500.2 | HK1 | CACATGGAGTCCGAGGTTTATG | CGTGAATCCCACAGGTAACTTC |
| 3099 | NM_000189.5 | HK2 | GAGCCACCACTCACCCTACT | CCAGGCATTCGGCAATGTG |
| 54704 | NM_001161779.2 | PDH | AGCGCCTCTCTAAAATGCTT | AGCCACTCACTATTCTTGGTT |
| 197257 | NM_153486.4 | LDHD | ATGGCAACTCTAAAGGATCAGC | CCAACCCCAACAACTGTAATCT |
| 2752 | NM_001033044.4 | GLUL | TTCCGTAAGGACCCTAACAAGC | CTCACCATGTCCATTATCCGTT |

| | | | | |
|---|---|---|---|---|
| 1373 | NM_001122633.3 | CPS1 | TGCCAAAACCTACAAGATGTCC | CCCCTCATTTGTTTGATCGTTG |
| 80017 | NM_001102368.3 | DGLUCY | CGCAGACACCTAGATTTGACCAC | CCAGGAATCTTCTTCGCAGCAA |
| 8659 | NM_003748.4 | ALDH4A1 | CCACCCCTTCTTCATGCAC | TACCTCCCACATGGCCGAT |
| 27165 | NM_013267.4 | GLS2 | TTCCTGCGGCATGTATGACT | GACAGGCACATCATTCCCAT |
| 5471 | NM_002703.5 | PPAT | AAATTGTCAGCCCTTCGTTG | AGACCAATACCATGACGCAGA |
| 790 | NM_004341.5 | CAD | CCAGCTTTTGCCCATACCAG | CACCGTGACACAGTTGCCAT |
| 2673 | NM_001244710.2 | GFPT1 | TAGAACACACCAATCGCGTCA | TGATCTCCTGCAGTTCGTT |
| 2746 | NM_005271.5 | GLUD1 | TCAGCTATGGCCGTTTGACC | GTCTTTCTCAGATGCACCCGAT |
| 6999 | NM_005651.4 | TDO | TAGAGTCAAACCTCCGTGCTT | TTTGCTGGCTCTATTCACACC |
| 8564 | NM_003679.5 | KMO | TGCTGCTGAGAAATACCCCAA | ATCTGACAGTTGAATAGGCTCCA |
| 1543 | NM_000499.5 | CYP1A1 | ACCCCTGATGGTGCTATCGAC | ACGCTGAATTCCACCCGTTG |
| 23475 | NM_014298.6 | QPRT | GTGCCCAAAATCCACTAGTCCT | GCCACTGACCCTAAAGATGTGT |
| 23498 | NM_012205.3 | HAAO | CCCGGTCTGCAACAAGCTC | TTTCCCTTGCTCCAGGACTCG |
| 4128 | NM_000240.4 | MAOA | TCATTCTTGCCCGGAAAGCTG | AGAGTACTGCTCCTCACACCA |
| 4129 | NM_000898.5 | MAOB | ACTATGCTGCCATAATGGGA | ACCTTGGCATAGAGTTCACA |
| 169355 | NM_194294.5 | IDO2 | CTTGCCCTTCCATTTGTCGAA | CCAATTTCCAGGAATCCGTCT |
| 1644 | NM_001082971.2 | DDC | CTTATCACTGACTACCGGCAT | GGATATAAGCCTGCAGTCCTT |
| 196 | NM_001621.5 | AhR | TCTTCAGCCACTTCATCATCCG | CATGTGAACTTGCTGACGTCCA |
| 125061 | NM_001010982.5 | AFMID | GTACTCCCAGTGCAGACCC | ACCAGGCTCCAAGTATCGTC |
| 8942 | NM_001199241.2 | KYNU | TTTTAAGCCTACGCCAAAACGA | CCTCTCTTGGCTTTATCATCCG |
| 316 | NM_001159.4 | AOX1 | TCAGCAAGTGCCCTAATGCAG | AAGGCTGACACAAATTCCC |
| 438 | NM_001171038.2 | ASMT | CAGCAAACTATCGCAAGGAC | ACATGCATTCCTTAGCCAGA |
| 51166 | NM_001286682.2 | AADAT | GCACCCCAGCACTTTTAACCA | AGCTGCCAGTATTGCATCCTT |
| 15 | NM_001166579.2 | AANAT | CAGGCTCTTCTCAGGCATCG | TCCCAGCACCTTCAACCTCC |
| 121278 | NM_173353.4 | TPH2 | AATATTACACCGGGATCCATGC | TCCCTTGTTGCCTTTGTCGTC |
| 5694 | NM_002798.3 | PSMB6 | CCTACCAGCTCGGTTTCCAC | TGATTCCCGCCATCAGGTCT |
| 11047 | NM_007002.4 | ADRM1 | TACATTCAGCAGACGGACGAC | AACTTCAGCACGTAGACCCT |
| 5686 | NM_002790.4 | PSMA5 | CCATGTTTCTTACCCGGTCT | TTCTCCACAGCTAGGCACAC |
| 5700 | NM_002802.3 | PSMC1 | CCCTGCCTGATGAAAAGACGAA | TTAGCCATGATCAGGTCGTCCA |
| 5710 | NM_001330692.2 | PSMD4 | ACCGACAAGGCAAGAATCACA | ACTTTCTCCTTCTTGAGGCGTTT |

**PS: All synthesized by Tsingke Biological Company**
